# The dominance of gene expression controlled by *trans*-eQTL hotspots contributes to phenotypic heterosis in maize

**DOI:** 10.1101/2025.11.03.686376

**Authors:** Gen Xu, Xuerong Yang, Mingyue Zhang, Congbin Kang, Ziang Tian, Yunhui Qi, Meijie Luo, Peng Liu, Jeffrey Ross-Ibarra, Jinliang Yang, Hongjun Liu

**Affiliations:** Department of Agronomy and Horticulture and Center for Plant Science Innovation, University of Nebraska-Lincoln, Lincoln, NE 68583, USA; State Key Laboratory of Wheat Improvement, College of Life Sciences, Shandong Agricultural University, Tai’an 271018, China; Department of Statistics, Iowa State University, Ames, IA 50011, USA; Beijing Key Laboratory of Maize DNA Fingerprinting and Molecular Breeding, Maize Research Institute, Beijing Academy of Agriculture and Forestry Sciences, Beijing 100097, China; Department of Evolution and Ecology, Center for Population Biology, and Genome Center, University of California, Davis, CA 95616, USA

## Abstract

Heterosis, or hybrid vigor, is a key phenomenon in genetics research and agricultural production, and has been primarily attributed to non-additive genetic effects such as dominance — a prevailing consensus shaped by decades of empirical research and theoretical debate. Although dominance may arguably arise from distal modifiers, their selective advantage is debated due to presumably small individual effects. To address this long-standing question, particularly how genetic dominance manifests at the transcriptomic level and contributes to phenotypic heterosis, we integrated transcriptomic and phenotypic data from a large population of maize hybrids and their inbred parents. We found that ∼ 30% of the expressed seedling genes in a significant proportion of hybrids exhibited expression patterns deviating from the average of the two parents, indicative of non-additivity. Further analysis suggests that while hybrid gene expression *per se* is primarily regulated by *cis*-eQTLs, expression dominance (or non-additivity) is disproportionately controlled by *trans*-eQTLs. These *trans*-eQTLs cluster into hotspots that regulate the non-additivity of hundreds of target genes, mostly within co-expression networks, and are notably enriched for transcription factors (TFs). Focusing on one such hotspot, we functionally validated a classical maize gene *ZmR1*, a basic helix-loop-helix (bHLH) TF associated with multiple seedling trait heterosis, as a candidate regulator of expression dominance across hundreds of genes. Overexpression of *ZmR1* enhances expression dominance of downstream genes and increases phenotypic heterosis in both seedling and adult traits. Further experiments confirmed its direct regulatory role in modulating genes involved in anthocyanin biosynthesis and lignin metabolism, driving transcriptome-level dominance. These results provide empirical support for the modifier hypothesis under an omnigenic model, suggesting that heterosis arises not from the modification of a single gene’s inheritance but through the coordinated regulation of hundreds of phenotype-associated genes, thereby helping to reconcile the long-standing debate over the genetic basis of dominance in heterosis.

## Introduction

Heterosis, or hybrid vigor, refers to the superior performance of hybrid progeny relative to their parents. Heterosis is widespread in cereal crops (Schnable and Springer, 2013), vegetables (Yu et al., 2021), and livestock (Dickerson, 1973), and has a profound impact on agricultural productivity. The genetic basis of heterosis has been debated for over a century, with three classical quantitative genetic models proposed: dominance (or “pseudo-overdominance”) (Davenport, 1908; Jones, 1917; Kusterer et al., 2007; Yu et al., 1997), overdominance (East, 1908; Shull, 1908), and epistasis (Yu et al., 1997). Supported by both theoretical and empirical evidence (Birchler et al., 2003; Charlesworth, 2018; Crow, 1998), the dominance hypothesis has recently regained attention, centering on the core idea of directional dominance — that the beneficial effects of dominance tend to act in the same direction (Lamkey and Edwards, 1999) — as well as the complementation of widely distributed and slightly deleterious mutations throughout the genome in subdivided populations (Liu et al., 2017; Yang et al., 2017).

The early theoretical frameworks for the mechanism of dominance laid the critical groundwork for understanding heterosis. Fisher proposed that dominance evolves through selection for “modifier” alleles that suppress the phenotypic effects of deleterious mutations (Fisher, 1928). Nonetheless, Wright (1929) mathematically demonstrated that modifiers that increase dominance, or masking of deleterious recessive alleles in hybrids, have only weak selective advantages and therefore are unlikely to be fixed rapidly under natural selection. Instead, Wright argued that dominance arises inherently from nonlinear biochemical interactions in metabolic pathways (Wright, 1934). Subsequent studies aligned with Wright’s model, demonstrating that dominance depends on the functional importance of genes and their position in metabolic networks (de Vienne et al., 2023; Kacser and Burns, 1981). These insights were later extended by Huber et al. (2018), who showed that dominance effects correlate with gene expression optimization and dosage sensitivity, suggesting that both evolutionary constraints and molecular mechanisms influence dominance and heterosis. Modern genomic approaches have expanded heterosis research to genome-wide expression studies, with a key focus on distinguishing between additive (parental mid-parent-like) (Guo et al., 2003, 2006; Stupar and Springer, 2006; Thiemann et al., 2014; Zhou et al., 2019) and non-additive (transgressive or dominant) patterns (Auger et al., 2005; Baldauf et al., 2016; Jahnke et al., 2010; Swanson-Wagner et al., 2009). However, most studies have been limited by small sample sizes, with the exception of a recent study using a QTL mapping population in maize (Pitz et al., 2025).

In this study, we investigated the genetic basis of dominance effects on gene expression and their contribution to phenotypic heterosis by generating population-wide RNA-seq data from a large number of hybrids and their inbred parents. Through integrative analysis, we identified population genetic patterns associated with gene expression dominance. In particular, we discovered an eQTL hotspot containing *ZmR1* locus, a bHLH transcription factor (TF) from the *R1* gene family (Tonelli et al., 1994), which regulates the expression dominance of hundreds of genes and is associated with phenotypic heterosis of several traits. Through detailed analysis of overexpression lines, we demonstrated that hybrids overexpressing *ZmR1* outperformed their wildtype counterparts. These results suggest that the identified TF *ZmR1* might serve as an example of Fisher’s “modifier” to regulate downstream gene expression, driving deviations from mid-parent values for hundreds of its regulated genes, which cumulatively contribute to phenotypic heterosis. The empirical evidence supports Fisher’s modifier model — not as a regulator of a single gene, but as a hotspot influencing the expression dominance of hundreds of downstream genes — placing it under selective advantage and potentially reconciling Wright’s argument.

## Results

### Gene expression dominance in maize seedlings from a blocked diallel population

We generated *n* =599 hybrids by crossing *n* =203 diverse inbred lines according to a blocked partial diallel design (**Materials and Methods, Figure 1A, S1**, and **Table S1**). The population structure analysis assigned most of these inbred lines to six previously defined heterotic groups (Li et al., 2022): Stiff Stalk (SS), Non-Stiff Stalk (NSS), Iodent, group A (PA), group B (PB), and Sipingtou (SPT) (**Figure S1A-C**). To minimize the confounding effects of genetic dosage and residual endosperm-derived nutrition during early seedling development (Jahnke et al., 2010), we dissected embryos for germination and seedling trait assessment (**Figure 1B-C** and **Figure S2A**). Consistent with previous studies (Paschold et al., 2010; Springer and Stupar, 2007; Yang et al., 2017), hybrids outperformed inbred parents across all seven seedling traits (**Figure S3**), with mid-parent heterosis (MPH) values ranging 12.6% – 81.2% for seedling traits (**Figure 1E, Table S2**), as compared to 10.8% – 145.8% for agronomic traits collected from replicated field trials using the same population (**Figure S2B, Table S2**, and **Materials and Methods**).

**Figure 1.**
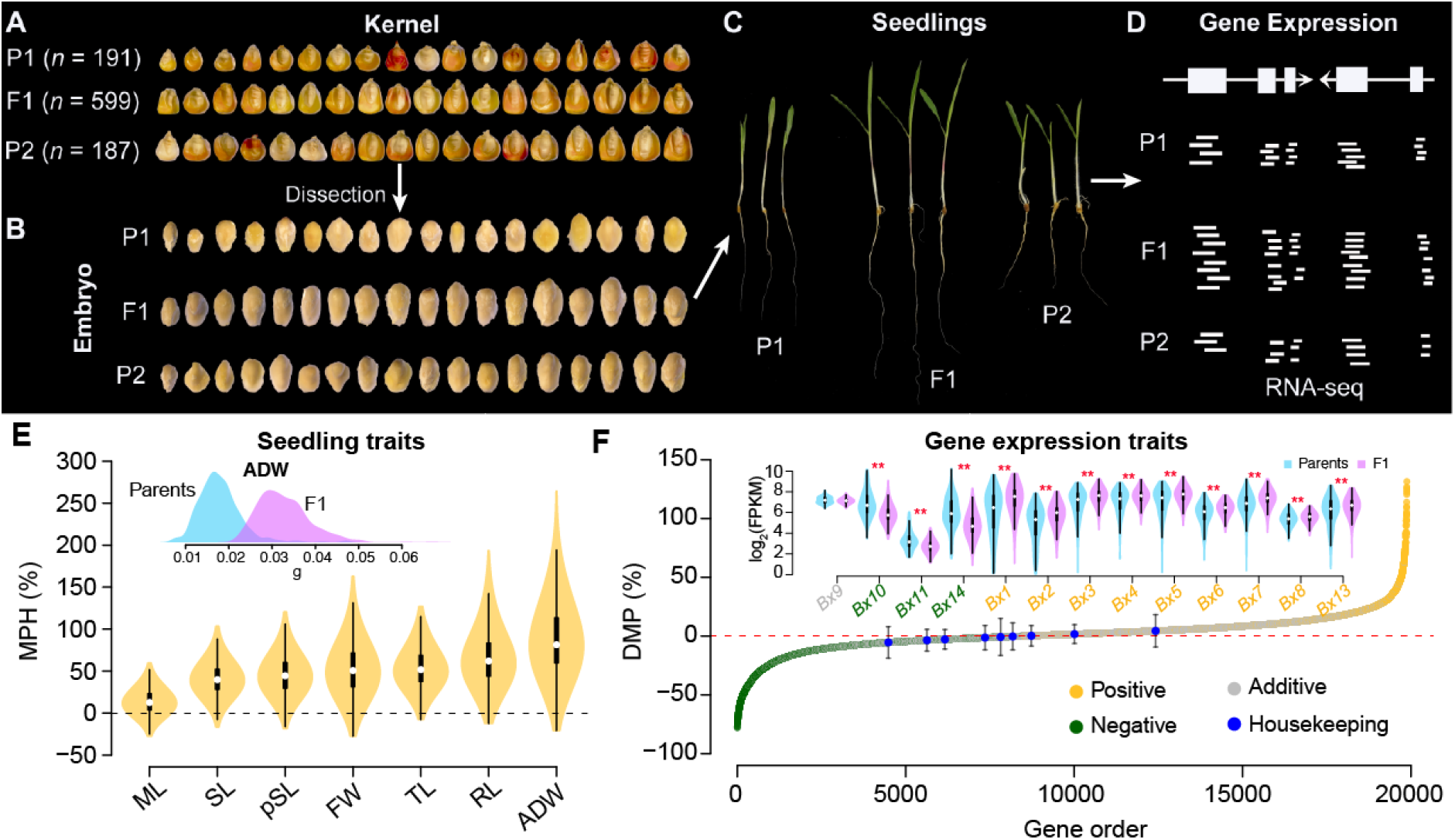
Schematic illustration of data collection and heterosis distributions. (**A**) Kernels were dissected to remove the endosperm, retaining only the embryo for seedling trait measurements and subsequent RNA-seq experiments. (**B-C**) Seedlings germinated from manually dissected embryos were used to collect seedling traits (**C**). (**D**) The two-week-old whole plant tissues were subjected to RNA-seq experiments. (**E**) Distribution of mid-parent heterosis (MPH) values for seedling traits. The inset shows the phenotypic distributions of hybrids and their inbred parents for the average dry weight. FW, fresh weight; ADW, average dry weight; ML, mesocotyl length; RL, root length; SL, seedling length; pSL, partial seedling length; TL, total length. (**F**) Deviation from mid-parent value (DMP) of gene expression in hybrids. Genes are categorized into three groups: positive (yellow), negative (green), and additive (gray). Blue dots represent DMP values from housekeeping genes, and error bars indicate standard deviation. In the upper panel, the genes involved in the benzoxazinoid (Bx) metabolic pathway, with yellow and green colors indicating positive and negative DMP, respectively. Asterisks indicate significance levels between hybrids and parental lines, as determined by the Wilcoxon test (**, *P* ≤ 0.01).

Using two-week-old whole-seedling tissues, we performed transcriptome profiling of hybrids and their inbred parents (**Figure 1D, Table S3**, and **Materials and Methods**). To ensure comparability across genotypes, RNA-seq libraries were prepared with standardized tissue quantities (or equivalent RNA input). Following RNA-seq data processing (see **Materials and Methods**), we identified *n* =26, 020 and *n* =27, 316 actively expressed genes (or median FPKM *>* 1) in the inbred and hybrid populations, respectively. We found a set of conserved housekeeping genes (Lin et al., 2014) exhibiting stable expression patterns in both inbreds and hybrids (**Figure 1F** and **S3**), suggesting the feasibility of direct gene-level comparisons between hybrids and their parental lines. Consistent with findings in other systems (Baldauf et al., 2022), most hybrids (85.3%, 511/599) exhibited more expressed genes than either parent, indicating a general increase in overall transcriptional activity in hybrids (**Figure S4**). Among *n* =19, 880 actively expressed genes, their dominance levels exhibit a broad distribution (**Figure 1F**), with a total of 6,458 genes (32.5%) from 599 hybrids significantly deviated from additivity (Kolmogorov-Smirnov test, FDR =0.05, see **Materials and Methods**), including 3,311 (16.6%) with positive and 3,147 (15.8%) with negative deviation from mid-parent values (DMP) (**Table S4**). For example, genes within the benzoxazinoid (*Bx*) pathway — a well-characterized and biologically coherent secondary-metabolite pathway (Wang et al., 2018), exhibit variation in the sign and direction of heterosis, likely due to their position and function within the pathway (**Figure 1F** and **S5**).

### Genetic and evolutionary determinants of gene expression patterns

Genes exhibiting extreme expression levels in maize inbreds are likely dysregulated due to the accumulation of deleterious mutations (Kremling et al., 2018). To investigate this, we quantified the derived deleterious genetic loads in inbred lines and found that, indeed, genes with extreme expression levels in the inbred parental lines showed elevated mutational burdens in their upstream regulatory regions (**Figure 2A** and **Materials and Methods**). Genes under stronger purifying selection are enriched among maize-sorghum syntenic orthologs and are significantly overrepresented in non-additive genes (permutation *P*-values =0.001, **Figure S6**). Moreover, evolvability, measured as coefficients of gene expression variation (Garfield et al., 2013), showed strong correlations with DMP values, suggesting that genes possessing greater evolvability maybe more responsive to hybridization and functional innovation (**Figure 2B**). Notably, genes exhibiting high levels of dominance showed increased SNP-based heritability 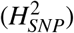 in hybrids when considering both additive and dominance SNP effects (**Materials and Methods**), indicating detectable and predictable genetic effects, typically with an oligogenic model (i.e., upstream regulatory variation). In contrast, genes showing high evolvability (i.e., high capacity for expression change over generations) displayed reduced heritability, reflecting a highly polygenic basis in which many small-effect loci contribute to changes in expression dominance (**Figure 2C**).

**Figure 2.**
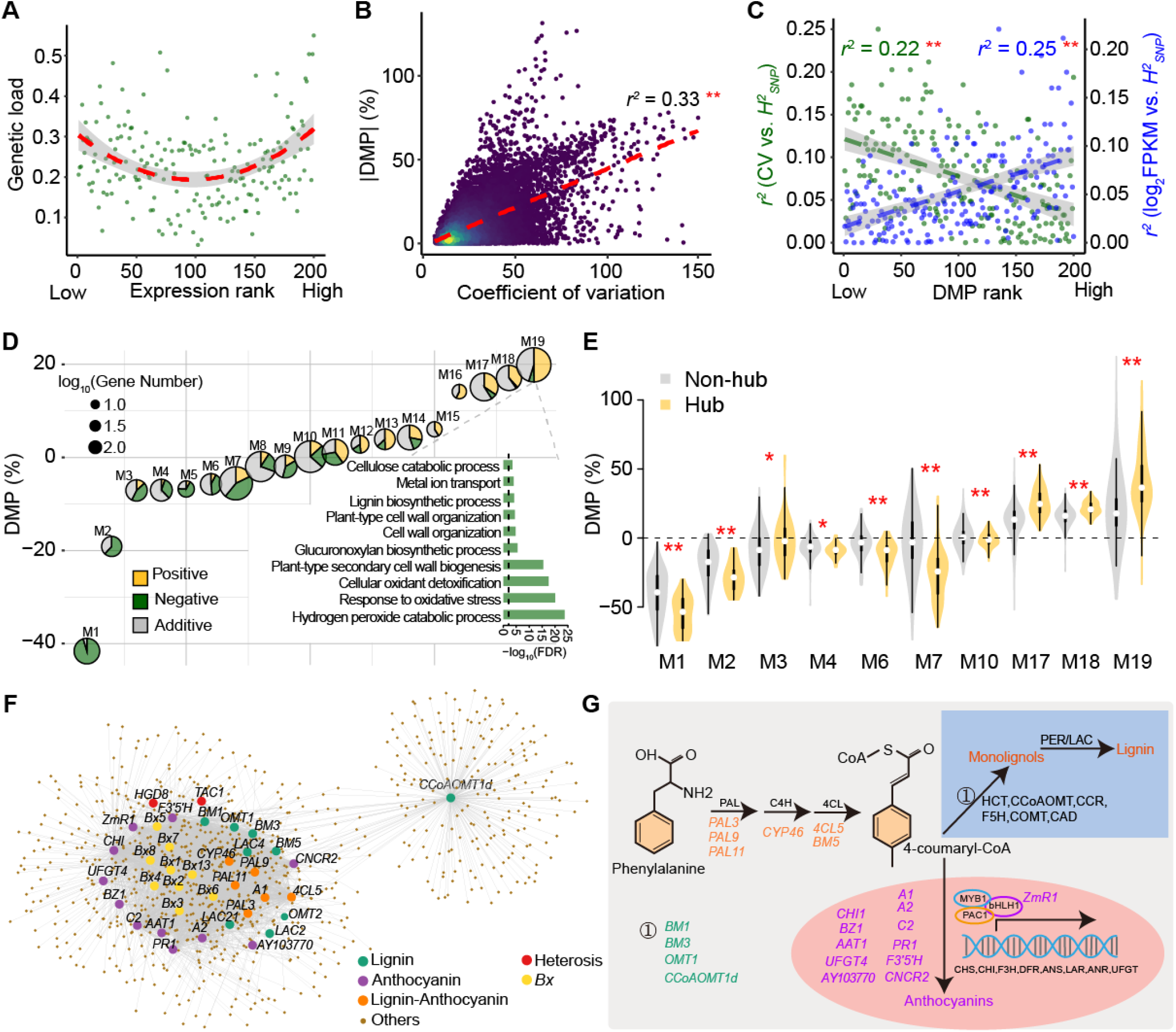
Genomic features and gene expression networks associated with expression dominance. (**A**) The median genetic load within the 5-kb upstream regions of genes using inbreds. Genes were stratified into 200 equal groups based on their expression levels. (**B**) Correlation analysis between the absolute deviation from mid-parent value (DMP) of gene expression and evolvability (coefficient of variation). (**C**) Correlation analysis between the absolute DMP of gene expression and two heritability measures: (1) heritability of evolvability, represented by the correlation between the coefficient of variation and SNP-based heritability; and (2) heritability of expression levels in hybrids, represented by the correlation between log_2_-transformed expression levels and SNP-based heritability. In (**A-C**), the dashed lines define the best-fit regression line. Red asterisks indicate the significant correlation at the *P* = 0.01 level for a Pearson correlation test. (**D**) Modules identified through co-expression analysis in hybrids. The inset barplot represents the significant GO terms in the biological process using genes from Module 19 (M19). (**E**) Gene expression DMP for the hub and non-hub genes of selected modules. Asterisks indicate significance levels based on Wilcoxon’s test (*, *P* ≤ 0.05; **, *P* ≤ 0.01). (**F**) A subset of the co-expression network of M19. Nodes (dots) vary in size and color, representing different gene categories. (**G**) Lignin and anthocyanin pathways. Genes marked with different colors represent distinct gene categories.

Furthermore, we constructed coexpression networks using hybrid gene expression data and identified 19 distinct gene modules (**Figure 2D, Materials and Methods**). Within each module, the hub genes show more pronounced dominance compared to the non-hub genes (**Figure 2E**). The largest module (Module 19, comprising *n* =1, 813 co-expressed genes) predominantly consists of genes exhibiting positive DMP and is enriched for genes involved in the hydrogen peroxide catabolic process or genes in the lignin metabolic pathway (**Figure 2D**, see GO term enrichment results for other modules in **Table S5**). Interestingly, Module 19 contains a number of anthocyanin genes (*n* =17, **Table S6**), which are a part of the lignin pathway (**Figure 2G**), and genes in the benzoxazinoid (*n* =9) pathway (**Figure 1F**), as well as homologous genes previously reported to contribute to heterosis in rice (**Figure 2F**), such as *TAC1* and *GHD8* (Huang et al., 2016).

### *Trans*-eQTL disproportionately regulates the expression dominance

To assess the genetic basis of expression dominance, we used DMP values to perform an eQTL analysis (Liu et al., 2020; Xiao et al., 2021). As a result, we identified *n* =14, 477 eQTLs associated with expression dominance, as well as *n* =19, 079 and *n* =12, 942 eQTLs for gene expression *per se* in hybrid and inbred populations, respectively (**Materials and Methods, Table S7**). Following a previous study (Gong et al., 2018), we defined eQTLs located within a 1-Mb flanking region of the target gene as *cis*-eQTLs. Those located more than 1 Mb away on the same chromosome were classified as *trans1*, and those located on different chromosomes were classified as *trans2*, to distinguish long-distance regulatory effects such as those observed for *tb1* (Xu et al., 2020). For example, Starch-Branching Enzyme (*SBEIIa*), a gene involved in the amylopectin biosynthesis pathway (Gao et al., 1997), has a *cis*-eQTL along with eight *trans1*- and five *trans2*-eQTLs detected in the hybrid population (**Figure 3A**). We detected seven *trans1*- and 28 *trans2*-eQTLs associated with the DMP value of *SBEIIa* expression, including several TFs likely involved in the regulation of starch biosynthesis (**Table S8**). A further summary of the eQTL analysis revealed that *cis*-eQTLs associated with expression *per se* were identified for over 70% of genes in both inbred lines (79.7%) and hybrids (77.3%), with most located within 5-kb upstream or downstream of the corresponding gene. In contrast, only 2.4% of genes with significant eQTL for DMP exhibited associations in *cis*, while 89.5% were located on chromosomes different from those of their associated genes, suggesting that the expression dominance is more likely to be regulated in *trans2*. Both *cis*- and *trans*-eQTLs are enriched in histone modification peaks (**Figure 3C**), suggesting that genomic variants reside in regulatory regions with the potential to influence transcription.

**Figure 3.**
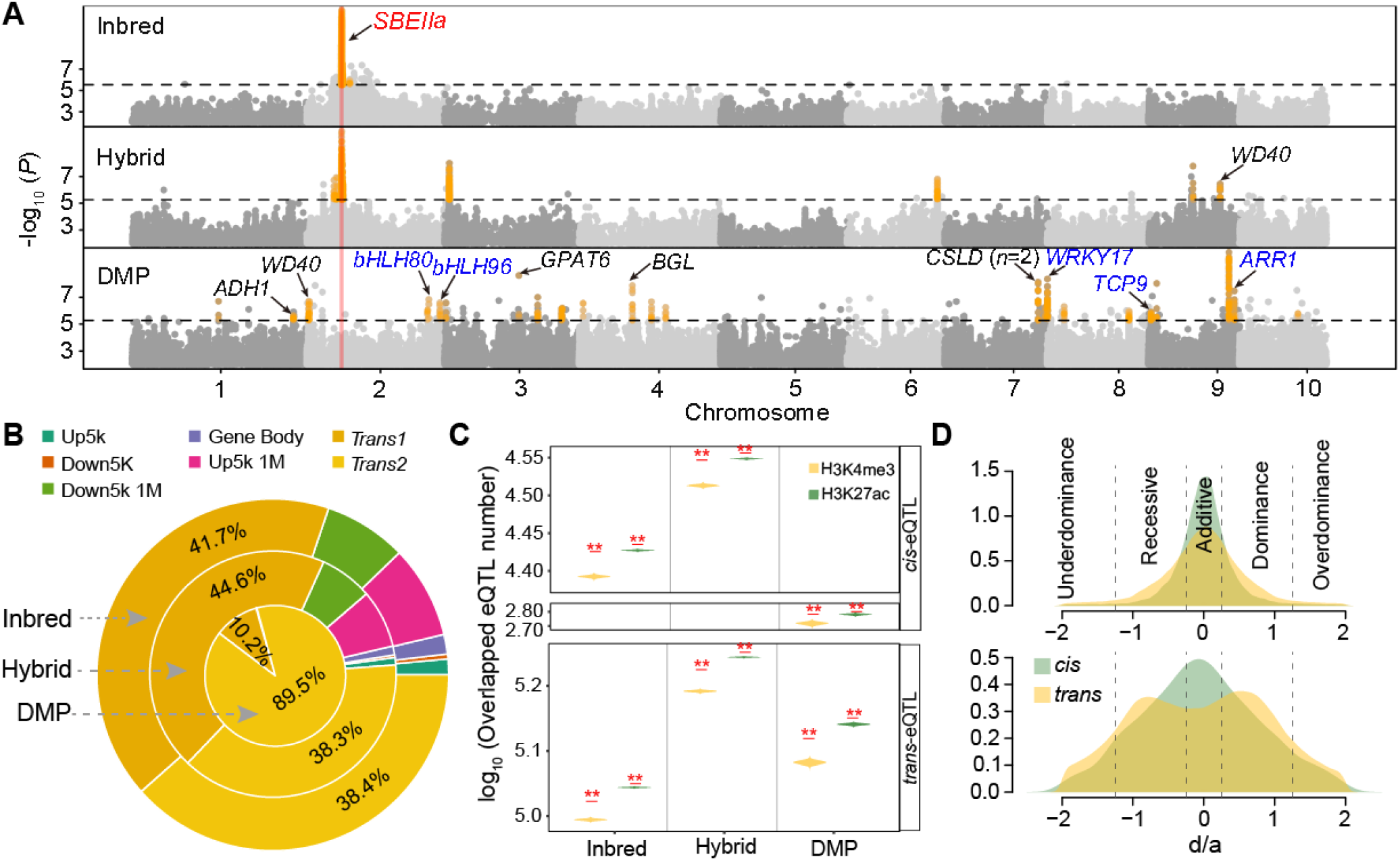
The *cis*- and *trans*-eQTLs for gene expression DMP. (**A**) Manhattan plots of GWAS for gene *SBEIIa* expression level in inbred, hybrid populations and its DMP. Genes with blue colour are transcription factors. (**B**) Proportions of eQTLs located in each genomic feature. *Trans1* refers to *trans*-eQTLs located on the same chromosome as their associated genes, while *Trans2* refers to those located on different chromosomes. (**C**) Comparison between eQTLs and histone modification peaks in the B73 genotype. The red asterisks indicate that eQTLs are significantly present in the histone modification peaks. Two asterisks denote *P* value < 0.01 from one-sided permutation test. (**D**) Dominance effects (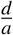 value) of eQTLs for gene expression traits *per se* in hybrids (up panel) and the transformed DMP traits (down panel). Orange and green lines indicate results from *cis* and *trans*-eQTLs.

**Figure 4.**
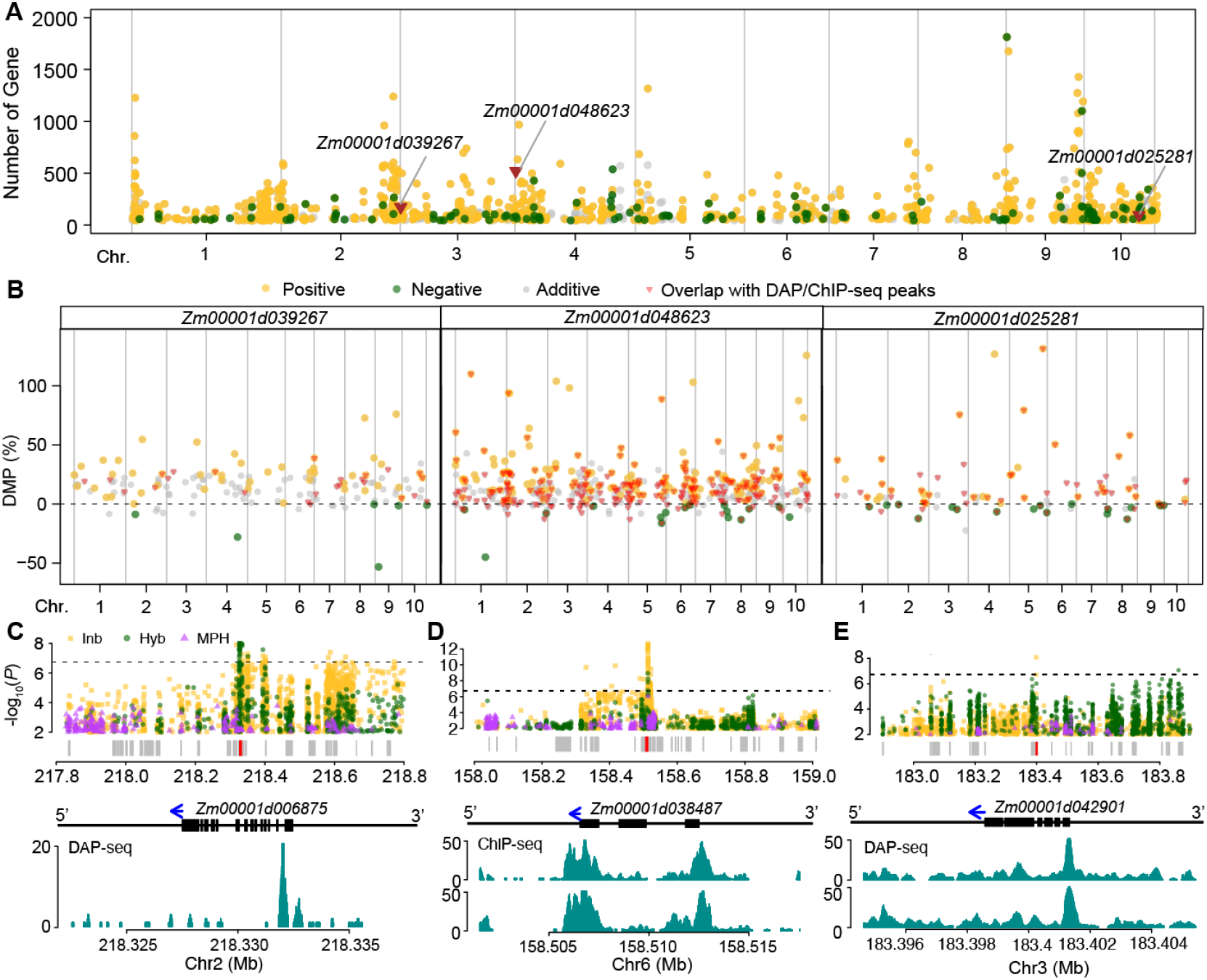
eQTL hotspots for gene expression dominance. (**A**) Distribution of the target gene number of eQTL hotspots. The red triangles represent the physical position of three candidate genes (*Zm00001d039267, Zm00001d048623*, and *Zm00001d025281*) under eQTL hotspots. The yellow and green dots indicate the target genes of eQTL hotspots enriched in positive and negative non-additive genes, respectively. (**B**) DMP values of the target genes of the three highlighted genes across the genome. (**C-E**) Examples show three hotspots where target genes overlap with DAP-seq and ChIP-seq peaks. The upper panels show the GWAS signals around the target genes for expression *per se* and expression DMP. The dashed horizontal lines indicate the GWAS threshold. The red and gray boxes indicate the target and other gene models. The middle panels show the target gene structure. The blue arrows indicate gene orientation. The bottom panels show the DAP-seq (**C, E**) and ChIP-seq (**D**) signals (green color) of the *trans*-eQTL candidate TF for each target gene.

To evaluate the genetic effects of eQTLs on expression, we further estimated the dominance index (the ratio of dominance to additive effects, *d/a*) for each eQTL detected in the hybrid population (**Figure 3D**) (see **Materials and Methods**). As expected for expression levels *per se*, both *cis*- and *trans*-eQTLs (including both *trans1* and *trans2*) showed dominance index values centered around 0, consistent with predominantly additive effects. In contrast, for DMP values, while *cis*-eQTLs remained centered near 0, *trans*-eQTLs displayed a bimodal distribution, with peaks indicative of incomplete recessive and incomplete dominance, supporting that *trans*-regulation drives deviation from additive expectation in gene expression dominance. Most of these deviations represent partial or incomplete dominance rather than strict complete dominance, consistent with the quantitative variation in the effects of gene dosage (Huber et al., 2018).

### *trans*-eQTLs cluster at specific genomic hotspots

In summarizing eQTLs, we found that eQTLs for gene expression *per se* are widely distributed throughout the genome, while eQTLs for DMP are concentrated in specific hotspots (**Figure S7**). We identified *n* =822 eQTL hotspots, each regulating the expression dominance of more than 45 genes (**Figure 3A, S8, Table S9**, and **Materials and Methods**). Of these, 675 (82%) eQTL hotspots are enriched for target genes with positive DMP, whereas only 92 (11%) are enriched for genes with negative DMP. Compared to the genomic regions under positive selection during maize breeding between the 1960s and the 1980s (Wang et al., 2020), we found that eQTL hotspots were significantly less likely to overlap with these regions than expected by chance (permutation *P* =0.01), consistent with the idea of heterozygote vigor.

Certain genes located within *trans*-eQTL hotspots, including those encoding TFs, may act as key regulators of transcriptional activity. To test this hypothesis, we retrieved maize TFs from the PlantTFDB database (Jin et al., 2016). Enrichment analysis revealed that TFs are significantly overrepresented within *trans*-eQTL hotspots (permutation *P* =0.004). Two TFs, *Zm00001d039267* (Auxin response factor 7) and *Zm00001d048623* (MYB-transcription factor 88), underlying two eQTL hotspots on chromosomes 3 and 4 (**Figure 3A**), have public DAP-seq (Galli et al., 2018) and ChIP-seq (Tu et al., 2020) data, separately. We processed the DAP/ChIP-seq data and compared the target genes of the two hotspots with the binding sites of the TFs. The analysis revealed that TF binding sites are significantly overrepresented among the target genes of *trans*-eQTL hotspots — by 60% and 40% more than expected by chance for *Zm00001d039267* and *Zm00001d048623*, respectively (permutation test, *P* =0.001, **Figure 3B-D**). Furthermore, we generated DAP-seq data for another TF gene, *Zm00001d025281*, annotated as AP2/EREBP, underlying a *trans*-eQTL associated with expression dominance for 96 genes, 67 (69.8%) of which overlapped with DAP-seq peaks (permutation *P* =0.001, **Figure 3B,E**).

### Gene expression contributes to seedling trait heterosis

To further assess the impact of gene expression on phenotypic heterosis, we conducted transcriptome-wide association studies (TWAS) for seedling trait heterosis (**Materials and Methods**). In total, 54 genes were associated with seedling traits *per se* and 25 for seedling heterosis, with *n* =11 genes (44%) overlapping between the two sets (**Figure 5A**). A large proportion of these trait-associated genes, particularly those associated with heterosis traits, are TFs. For example, the MADS-box gene *TU1*, previously implicated in the regulation of grain yield and leaf number above the ear (Li et al., 2024), was associated with both seedling fresh weight and its heterosis (**Figure 5A**, see other genes and their potential functions in **Table S10**). Another TF *ZmR1*, a bHLH that participates in the synthesis of anthocyanin pigmentation (Dellaporta et al., 1988; Luo et al., 2022), was strongly associated with one seedling trait *per se* (FW) and four seedling heterosis traits (FW, pSL, SL, and TL) (**Figure 5A**). The expression level of *ZmR1* is higher in most hybrids compared to their parental lines (**Figure 5B**), exhibiting a significant deviation from the mid-parent value (paired Student’s *t* test *P* = 2.9e-49, **Figure S9**). Furthermore, this gene is located within a *trans*-eQTL hotspot and potentially regulates the expression dominance of more than 230 genes, which are predominantly positive dominance genes (**Figure 5C**). We then investigated whether these targets are correlated with phenotypic performance. Indeed, correlation analyses revealed that the expression levels of most *ZmR1* target genes were positively correlated with the four seedling traits, and that their expression dominance (i.e., DMP) values were likewise positively correlated with the heterosis of the seedling trait (**Figure S10**). Notably, several of these targets are previously characterized genes, including *FHT1*, involved in the anthocyanin biosynthesis pathway (Wang et al., 2025); *CESA2*, annotated as a cellulose synthase (Zhang et al., 2024); and two photosystem-related genes — *CAB3* (Osadchuk et al., 2025) and *LHCB1* (Li et al., 2025). Consistent with the regulatory role of the eQTL hotspot, genes co-expressed with this candidate gene exhibit even stronger dominance (**Figure 5C**). Co-expression network analysis further reveals that the target genes of this eQTL hotspot are distributed across multiple modules, particularly in module 19, where they are co-expressed with a set of genes involved in lignin, anthocyanin, and benzoxazinoid biosynthetic pathways (**Figure 5D**). To validate the downstream targets of *ZmR1*, we randomly selected five potential target genes for a dual luciferase assay (**Materials and Methods**). The results demonstrated that *ZmR1* functions as a transcriptional activator for *Zm00001d021599, Zm00001d013133*, and *LHCB3*, while it exerts transcriptional repression on *Zm00001d042730* and *Zm00001d005478* (**Figure 5E**).

**Figure 5.**
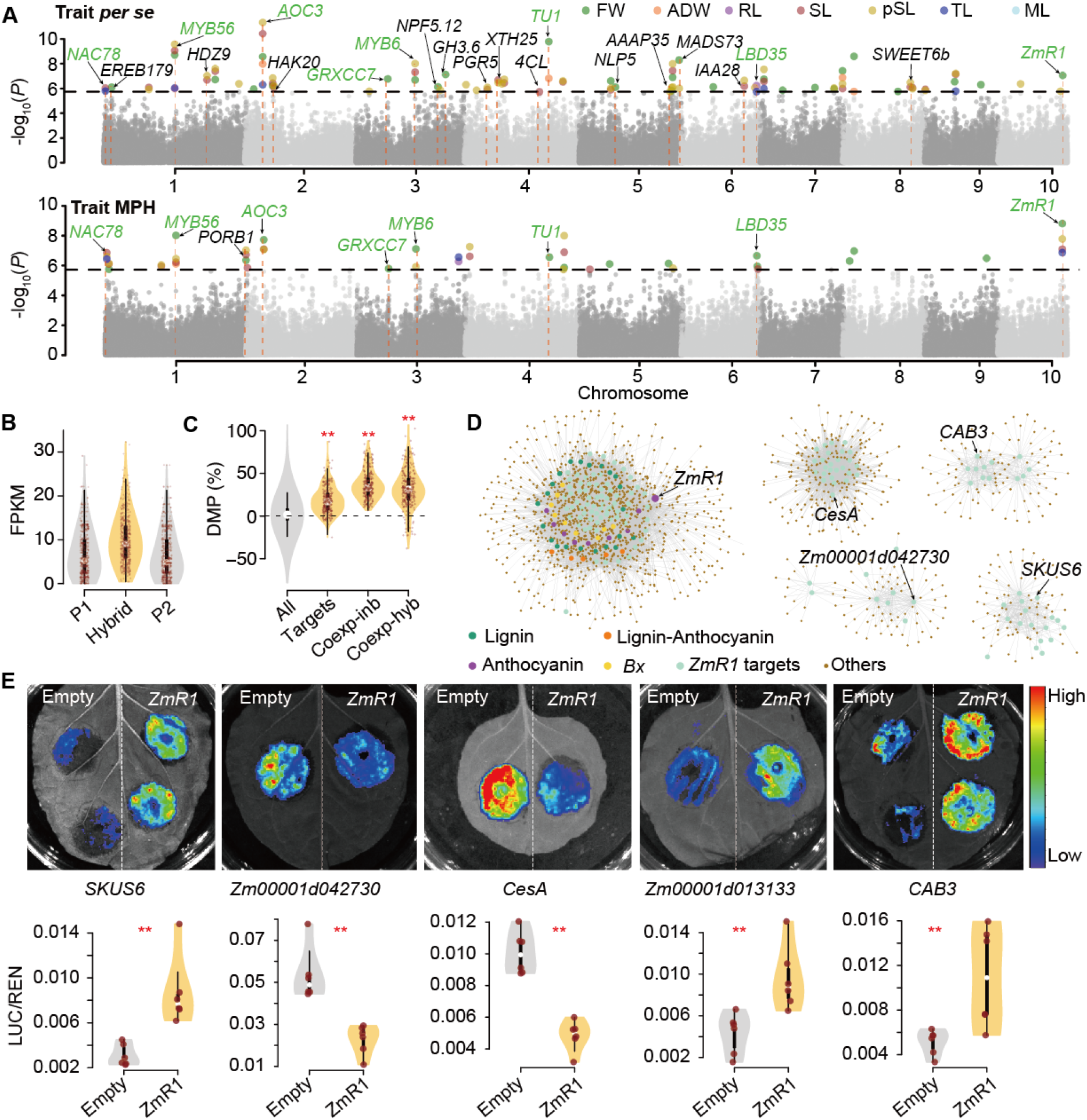
*ZmR1*, a candidate gene within the *trans*-eQTL hotspot, is associated with seedling trait heterosis. **(A)** Transcriptome-wide association studies (TWAS) for seedling traits *per se* and their heterosis. Gene names highlighted in green and red are associated with both traits *per se* and heterosis. (**B**) Expression level of *ZmR1* in hybrids and their parental lines. (**C**) Expression MPH of genes regulated by or co-expressed with *ZmR1* in inbreds and hybrids. (**D**) Co-expression network constructed from genes co-expressed with the targets of *ZmR1*. Node size and color indicate different gene categories. (**E**) Validation of *ZmR1*–target gene interactions using a dual-luciferase assay. For each target, the left image shows the empty vector control; the right shows co-expression with *ZmR1*. Asterisks indicate significance levels based on Wilcoxon’s test (**, *P* ≤ 0.01?.

### Overexpression lines of *ZmR1* influence trait heterosis

To quantify the phenotypic effects of *ZmR1*, we obtained three overexpression (OE) lines in the Jing724 background (Luo et al., 2022). We generated hybrids by crossing an elite inbred line, Jing92, with either the wildtype Jing724 and three OE lines. Both the OE lines and their hybrids, as previously reported (Luo et al., 2022), displayed a distinct purple coloration in the seedling leaves (**Figure 6A**). To evaluate phenotypic outcomes, we measured four seedling traits in a replicated greenhouse experiment (**Materials and Methods**). Overall, the OE allele in Jing724 reduced the trait values in inbred lines but increased them in hybrids (**Figure S11A**). Furthermore, OE hybrids exhibited progressively enhanced heterosis for fresh weight, seedling length, and root length (**Figure 6B**), corresponding to increasing levels of *ZmR1* expression (**Figure S12**). Interestingly, results from replicated field experiments indicated that OE hybrids exhibited significant increases in the trait *per se* (**Figure S11B**), along with enhanced heterosis for plant height and stalk stiffness (**Figure 6B** and **S13**).

**Figure 6.**
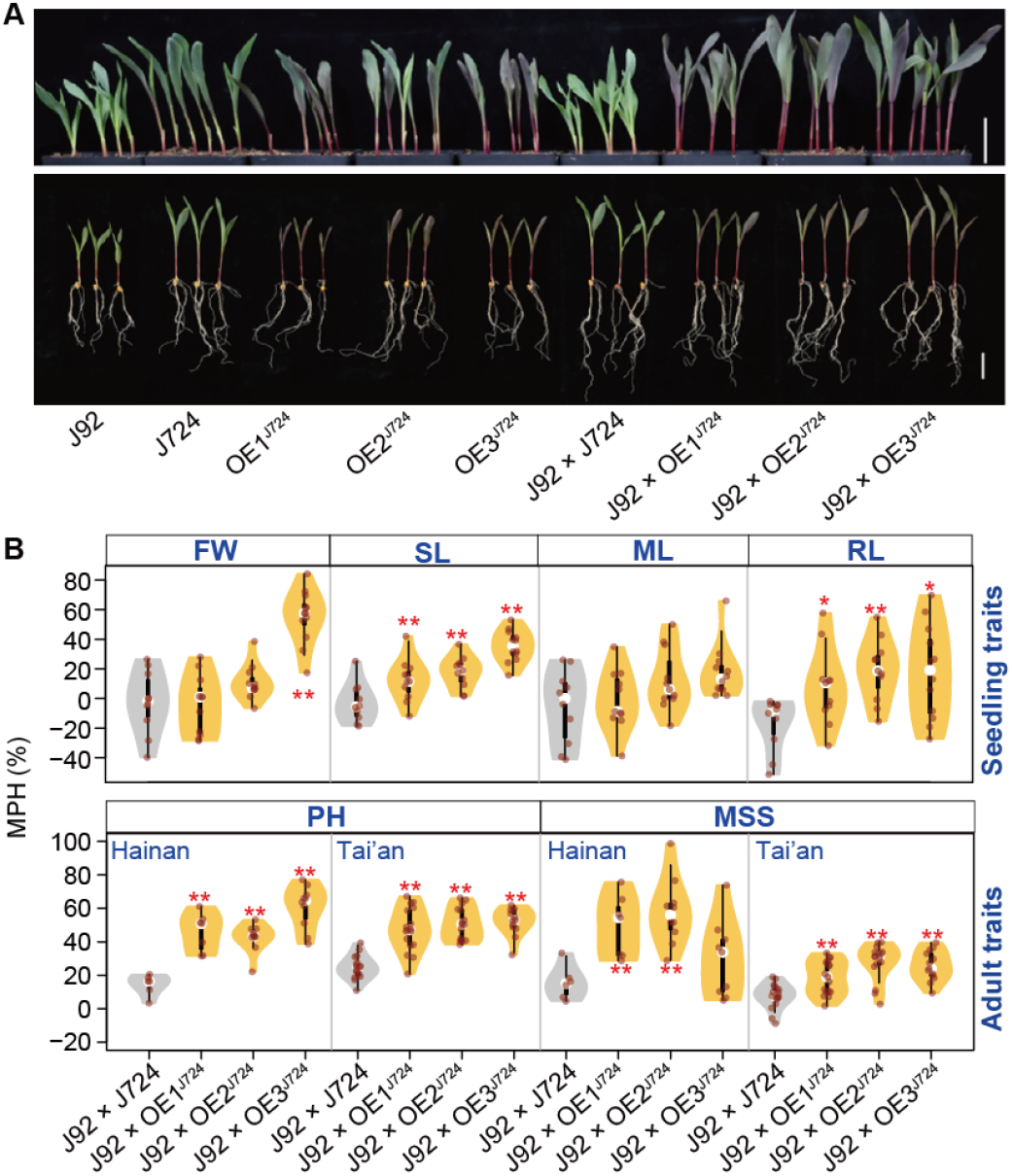
*ZmR1* affects seedling and adult trait heterosis. (**A**) The overexpression (OE) lines of *ZmR1* and their hybrids. Scale bar = 5 cm. (**B**) Seedling and adult trait heterosis in hybrids derived from crosses with OE lines. Adult traits were collected from two locations: Hainan and Tai’an. FW, fresh weight; SL, seedling length; ML, mesocotyl length; RL, root length; PH, plant height; MSS, middle stalk stiffness. Asterisks indicate significance levels between the OE lines and their corresponding wildtypes based on Wilcoxon’s test (*, *P* ≤ 0.05; **, *P* ≤ 0.01). The full names of inbred lines J724 and J92 are Jing724 and Jing92, respectively.

Since stalk stiffness is a critical trait associated with lodging resistance (Robertson et al., 2016), and lignin is a major contributor to cell wall rigidity (Liu et al., 2016), we further examined lignin accumulation in stem tissues. We performed phloroglucinol staining (**Materials and Methods**), which specifically detects lignin (Tang et al., 2013), and found that OE lines, particularly OE hybrids, exhibited more intense staining, indicating elevated lignin content (**Figure 7A**). These visual observations were consistent with quantitative measurements of lignin content, which showed significantly higher levels in the OE hybrids relative to their matched wildtypes (**Figure 7B**). Furthermore, MPH for lignin content was significantly increased in hybrids derived from OE lines (**Figure 7C**).

**Figure 7.**
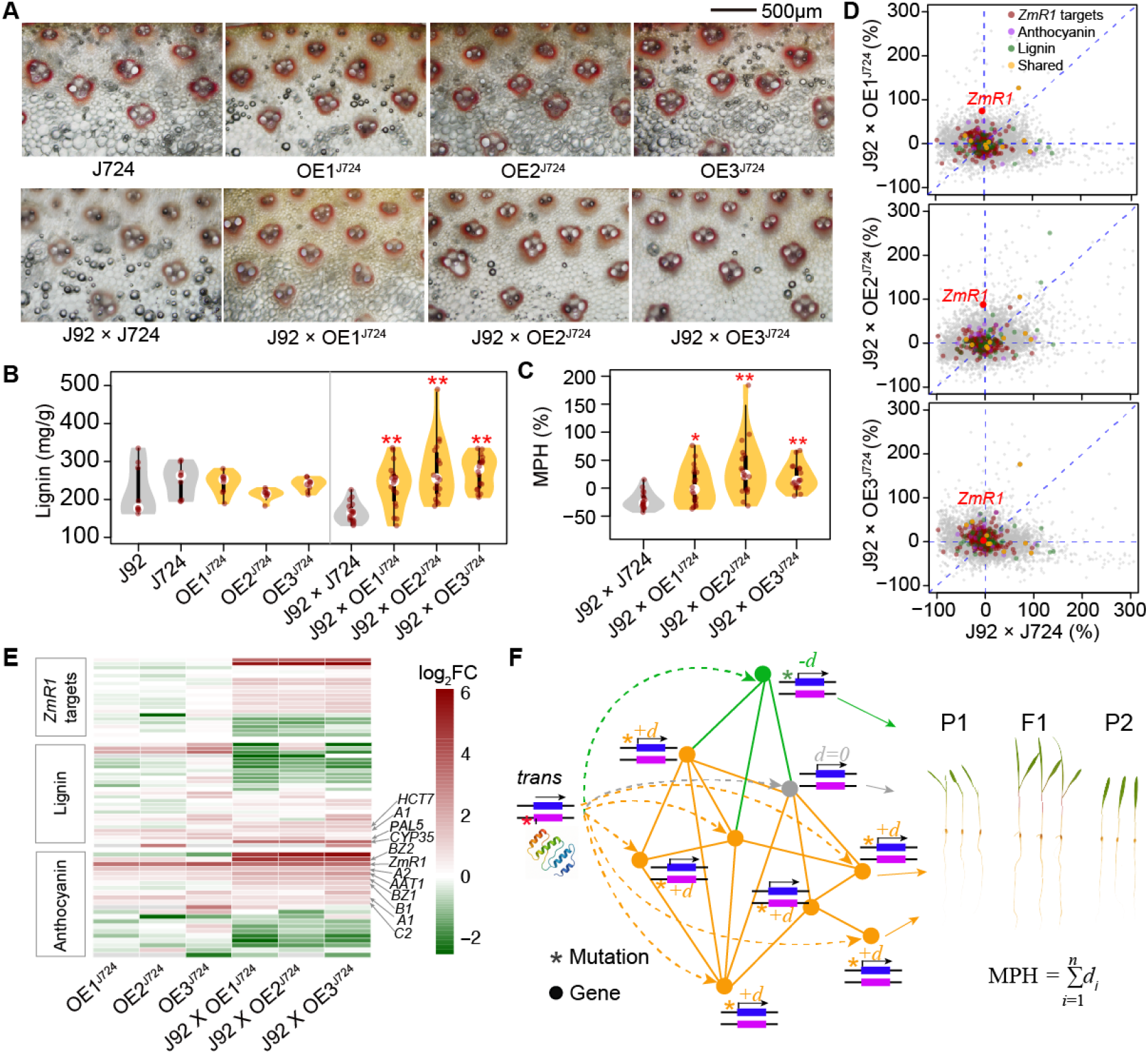
Overexpression of *ZmR1* enhances lignin accumulation and expression dominance. (**A**) Phloroglucinol staining of stem cross-sections from *ZmR1* OE lines. Scale bar = 500 µm. (**B–C**) Lignin content (**B**) and its heterosis (**C**) in *ZmR1* OE lines. Asterisks indicate significance levels between OE lines and their corresponding wildtypes based on Wilcoxon’s test (*, *P* ≤ 0.05; **, *P* ≤ 0.01). (**D**) Comparison of expression dominance between hybrids generated from *ZmR1* OE lines and the corresponding wildtype hybrids. Dark red dots represent *ZmR1* target genes showing a non-additivity pattern, and orange dots indicate genes involved in the same pathway as *ZmR1*. (**E**) Log_2_-transformed expression fold changes (log_2_FC) between OE lines and their matched wildtypes for *ZmR1* dominance eQTL target genes and genes involved in lignin and anthocyanin biosynthetic pathways. (**I**) A proposed model illustrating the elevated dominance of gene expression. A *trans*-eQTL can regulate the expression of its target gene and downstream genes in the hybrid. “*” represents the alternative allele introduced by mutation.

In particular, at the transcriptome level, most of the target genes of *ZmR1* for the eQTL hotspot exhibited increased DMP values of these targets in OE hybrids (**Figure 7D**). We further assessed the expression dominance of genes co-expressed with *ZmR1* and found a similar pattern, with most genes showing an increased dominance in OE lines, both in inbreds (**Figure S14A**) and hybrids (**Figure S14B**). In addition to *ZmR1*, several key genes in the lignin and anthocyanin biosynthesis pathways, such as *HCT7, PAL5, A1, A2, BZ1*, and *AAT1*, also showed elevated expression *per se* (**Figure 7E**) and increased expression dominance (**Figure 7D**) in OE hybrids. Taken together, our results suggest that *ZmR1* enhances the dominance of hundreds of genes at the transcriptomic level, leading to phenotypic heterosis in both seedling and adult traits, specifically for traits related to lignin.

## Discussion

In this study, we used MPH as a proxy for trait heterosis, as it can be modeled as the sum of dominance effects (*d*) weighted by squared allele frequency differences (*y*^2^) between two parental lines across genome-wide loci (Falconer, 1989). In this framework, *y* represents the difference in allele frequency between populations, that is, two different heterotic groups, Stiff Stalk and Non-Stiff Stalk (Grzybowski et al., 2023), and *y*^2^ equals one only when parents are fixed for alternate alleles. This model highlights that directional dominance, in which dominance effects consistently favor the same direction, is required for positive heterosis. Consequently, the amount of heterosis is specific to each individual cross for a given trait (Lamkey and Edwards, 1999), depending on the pattern of allele divergence and dominance interactions throughout the genome, consistent with our observations for the heterosis of both seedling and adult traits, as well as previously published results (Hochholdinger and Baldauf, 2018; Yang et al., 2017). In this context, the ‘dominance’ model we refer to represents directional dominance at the population level, where most genes show partial or incomplete dominance in expression rather than complete complementation of recessive alleles. Similarly, when considering gene expression as a trait, our results indicated that approximately 30% of expressed seedling genes (*n* =6,458) exhibited expression dominance in *>* 5.1% of hybrids, while only about 500 genes showed dominance in over 300 hybrids (50%) — suggesting that the majority of maize genes exhibit variation in expression dominance, whereas most genes in the majority hybrid families consistently follow an additive expression pattern (Pitz et al., 2025).

If we further hypothesize the phenotype as a function of genome-wide gene expression variation under an omnigenic inheritance model (Liu et al., 2019), phenotypic heterosis should reflect the cumulative effects of expression-level dominance acting in the same direction. This idea aligns with the observation that co-expressed genes, possibly involved in the same metabolic pathways (e.g., the *Bx* or lignin pathways), tend to exhibit coordinated patterns of positive or negative dominance. We found that expression dominance is largely associated with deleterious genetic load in upstream regions and is therefore heritable. However, *cis*-regulation alone, while accounting for the majority detectable eQTLs for the trait *per se* in hybrids, rarely causes gene expression to deviate from additive inheritance patterns, and its ability to evolve over generations appears limited. In fact, despite decades of improvement in yield, the magnitude of heterosis in maize has remained largely constant (Duvick, 1999; Troyer and Wellin, 2009). Expression dominance, calculated as the difference between hybrid expression and the average expression of the two parents, is therefore unlikely to arise from variants within the gene or its immediate upstream region. This is consistent with our finding that disproportionately more *trans*-eQTLs were detected for expression dominance, and that expression dominance is largely controlled by heterozygous regions of the genome (Pitz et al., 2025). At the molecular level, *trans* factors may function as diffusible proteins — either individually or as part of a complex — that bind to non-deleterous alleles in *cis*, recruiting the complete gene transcription machinery (Cramer, 2019), thus masking the deleterious effects of *cis*-regulatory variants (**Figure 7I**). Consistent with the omnigenic model (Liu et al., 2019), the expression levels of *trans*-eQTLs (peripheral genes) do not necessarily have direct effects on phenotypic heterosis (or are difficult to detect due to relatively small effect size). Instead, they can indirectly affect heterosis by modifying the expression dominance of hundreds of core genes. Such *trans*-eQTL modification in specific genes would have limited fitness effects and likely would not evolve, but modifiers that affect hundreds or thousands of genes could confer a selective advantage (Monroe et al., 2022). In support of this idea, a key insight from our study suggests that expression dominance in hybrids is primarily governed by *trans*-acting regulatory loci clustered into distinct regulatory hotspots. In particular, many of these hotspots contain TF genes, suggesting that they act as hub genes or master regulators orchestrating coordinated expression deviations from additivity, predominantly in the same direction.

We perform functional validation of the candidate transcription factor *ZmR1* and its downstream target genes using molecular, reverse genetic, and transcriptomic analyzes. The *ZmR1* gene is located within a major *trans*-eQTL hotspot and regulates the dominance of over 230 downstream genes, many of which are part of co-expression networks, exhibit mostly positive dominance, and are involved in anthocyanin, lignin, and photosynthesis pathways. Functional validation through dual-luciferase assays and expression analysis in overexpression lines confirmed that *ZmR1* acts as a transcriptional activator or repressor for specific targets, and its activity enhances the dominance of transcriptome expression. It is important to note that *ZmR1* should not be regarded as a single “major heterotic gene”; rather, it serves as a representative example of a *trans*-acting regulatory factor that influences the expression dominance of hundreds of downstream genes. Its dosage-dependent effects exemplify how transcriptional regulators within *trans*-eQTL hotspots can collectively modulate phenotypic heterosis through partial dominance across the transcriptome.

Collectively, our results are consistent with the dominance (or incomplete/partial dominance) hypothesis contributing heterosis and provide empirical evidence that, at the gene expression level, regulatory factors — particularly TFs within *trans*-eQTL hotspots — may function analogously to the ‘modifiers’ proposed by Fisher, acting to buffer deleterious alleles in the *cis*-regulatory regions of their target genes. A master TF typically regulates a set of co-expressed genes in a manner of directional dominance, which can ultimately manifest as phenotypic heterosis in a genotype-specific fashion. It is important to note that heterosis in this context can be confounded with natural variation in developmental traits. In our model, the pathway in which the master regulator is involved plays a central role in driving the trait, making the observed heterosis inherently trait-specific. This proposed idea aligns with theoretical models of dominance rooted in biochemical nonlinearity and gene network buffering and provides a predictive framework for identifying and engineering regulatory elements to enhance heterosis in breeding programs.

## Materials and Methods

### Plant materials for blocked partial diallel mating design and field trial

We selected a diverse set of 203 maize inbred lines to generate hybrid crosses. The selected inbred lines represent a subset of the diverse lines of maize HapMap3 and are also part of the panel of global diversity of maize (Bukowski et al., 2018; Grzybowski et al., 2023). We grouped the inbred lines into 10 blocks based on their flowering time, and within each block, we made crosses following a partial diallel mating design, resulting in about 600 hybrids. Using these hybrids and their inbred parents, we conducted replicated field trial in multiple locations in China during the summer of 2021: Taian, Shandong (36°11’07”N, 117°08’05”E); Suihua, Heilongjiang (SH, 46°39’32”N, 126°58’05”E); and Harbin, Heilongjiang (HRB, 45°51’38”N, 126°49’03”E).

### Field phenotypic data collection

We measured plant height and ear height from at least six plants per line after pollination. After harvest, we collected grain weight, moisture level, and hundred kernel weight using a seed counter. Grain weight was adjusted to 14% moisture, and average grain yield was calculated by dividing the total grain weight by the number of plants in that line.

### Procedure for embryo dissection

We sterilized the seeds with 75% v/v ethyl alcohol for one minute, rinsed three times with sterile water, then with 10% NaClO for 10 minutes, and rinsed with sterile water three times. The seeds were then stored in sterile seed bags at 4 ° C for further treatment. Subsequently, the sterile seeds were soaked in sterile water overnight for 18 hours and then used to dissect embryos from other components of the seed, including endosperm and seed coat. The dissected embryos were then sterilized with NaClO 7% v / v and rinsed three times with sterile water. Every 12 embryos of each hybrid line were covered in sterile brown paper and hydroponically cultured in Hoagland’s nutrient solution (900 ml) in each container, following a previous study (Luo et al., 2021), and 24 embryos (every 12 embryos in 900 ml of nutrient solution) for each inbred line due to the lower germination rate. The embryos were cultured in a greenhouse at 50% relative humidity in an 18 h light / 6 h dark photoperiod. To ensure consistent growth conditions, sterile water was added to up to 900 ml of each container daily for 14 days. Then, at 8:00 am on day 14, the individual plant was rinsed with sterile water and dried with sterile filter paper. Finally, lines with at least four growth-consistent individuals were chosen for data collection. Three individual plants from each line were snap-frozen with liquid nitrogen and then stored at -80 °C for RNA extraction. Otherwise, the remaining individual plants were used to measure root and seedling traits.

### RNA preparation and RNA-seq library construction

Total RNA was extracted and pooled from three individual plants for each line using TRIzol ReagentTM (Life Technologies Corporation, Carlsbad, CA, cat#15596026) according to the manufacturer’s protocol. Then, total RNA was qualified and quantified with a cutoff of 28S/18S > 1.0 and RNA Integrity Number (RIN) 6.0 by using Agilent 2100 Bioanalyzer (Agilent, CA, USA). Subsequently, the qulified and quantified RNA was used for RNA-seq library construction. Libraries were prepared according to Illumina Standard total RNA-seq library preparation kit. Briefly, total RNA was purified and fragmented, then was used to synthesize first strand and second strand cDNA; after purification, the second strand cDNA was ligated with adapters. After further purification, the DNA fragments with 250 - 300 bp were enriched and purified as RNA-seq libraries. Finally, libraries were sequenced to generate 150-bp paired-end reads (PE150) using the DNBseq platform from BGI-Shenzhen.

### Gene expression quantification of RNA-seq data

Reads were first filtered using fastp 0.20.0 (Chen et al., 2018) with parameters to remove low-quality reads. All filtered reads were then mapped to the maize B73 AGP v4 reference genome (Jiao et al., 2017) using the aligner STAR 2.7.9a (Dobin et al., 2013) with default parameters. The bam files of uniquely mapped reads were then counted from raw reads assigned to reference gene models using featureCounts v2.0.2 (Liao et al., 2014). Raw read counts were then normalized using the TMM (trimmed mean of M values) normalization method implemented in edgeR (Robinson et al., 2010) to give CPM (counts per million) values for each gene. CPM values were then normalized by gene length to give FPKM (fragments per kilobase of exon per million reads) values. Genes with FPKM values < 0.1 in all samples were excluded, and those with FPKM > 1 were regarded as actively expressed, as described previously (Zhou et al., 2019).

### Calculation of mid-parent heterosis (MPH) for phenotypic traits and deviation from the mid-parent value (DMP) for gene expression

The middle-parent heterosis (MPH) for each phenotypic trait was calculated using the formula MPH = (F1 - mp) / mp, where F1 represents the phenotypic value of the hybrid and mp is the average value derived from the two parental lines. Similarly, for gene expression, the deviation from the mid-parent value (DMP) was calculated using the same formula as MPH. Only genes with FPKM values greater than 1 in both parental lines and their corresponding hybrids were included in the analysis. Additionally, each gene’s DMP value was retained only if the missing rate across the entire population was below 30%. We chose to quantify gene expression dominance using mid-parent deviation (DMP) rather than high-parent deviation (DHP) because DMP provides a statistically balanced measure of deviation from additive inheritance.

To identify genes that exhibit DMP for each hybrid, we developed a two-stage likelihood ratio test (2sLRT) for unreplicated RNA-seq count data and applied the test to obtain a p-value for each gene in each hybrid. If a gene is additive across all families, we expect the *P*-values for this gene to follow a Uniform(0,1) distribution across families. A one-sided Kolmogorov-Smirnov (KS) test was applied to identify genes whose distribution of *P*-values was stochastically smaller than Uniform(0,1), corresponding to genes that tend to be non-additive across families. The Benjamini-Hochberg method was applied to *P*-values from the KS test to control the false discovery rate (FDR) at 0.05. The proportion of all hybrids for a gene to be detected non-additive at an FDR of 0.05 was also calculated, and this proportion is at most 5.18% among genes with an insignificant KS test. We declare a gene tends to be non-additive based on both FDR control of the KS test and the threshold of minimal proportion of non-additive families (5.18%). A gene is classified as a positive non-additive gene if the median value of its expression of DMP in the hybrid population is greater than zero; otherwise, it is considered a negative non-additive gene.

### expression QTL (eQTL) analysis

Whole-genome sequencing (WGS) SNPs of parental lines were obtained from the Hapmap 3.2.1 reference panel (Bukowski et al., 2018), which can be downloaded from https://www.panzea.org/genotypes. Then, the hybrid genotypes were inferred by CreateHybridGenotypesPlugin in TASSEL v5.0 (Bradbury et al., 2007). After filtering out SNPs with minor allele frequency (MAF) < 0.05, approximately 21 and 11 million SNPs were retained in the inbred and hybrid populations, respectively. We performed eQTL mapping for each gene in the inbred and hybrid populations, as well as expression MPH, using a mixed linear model implemented in GCTA software (Yang et al., 2011). To account for inferred confounders that influence expression variation, we applied the R package PEER (Stegle et al., 2012) to estimate the hidden factors using the gene expression matrix from inbred and hybrid populations, respectively. In the analysis, the hidden factors were used as covariates together with the first three principal components (PCs) calculated from genome-wide SNPs using PLINK 1.9 software (Chang et al., 2015). The kinship matrices were calculated using the centered IBS method implemented in TASSEL 5.2 (Bradbury et al., 2007). The log2-transformed FPKM values were used as phenotypes in inbred and hybrid populations. The threshold for the significant association SNPs was set to 2.9e-6 (1/n, *n* =339, 202 is the number of independent SNPs) for the inbred population and 5.4e-6 (1/n, *n* =183, 864) for the hybrid population and expression MPH. Here, the independent SNP number was determined by using PLINK 1.9 with the indep-pairwise option, as described in previous studies (Rodene et al., 2022; Yang et al., 2022) (window size 100, step size 10, *r*^2^ ≥ 0.1).

We define eQTL peaks as those identified by considering a 50 kb window upstream and downstream of significant SNPs. Overlapping regions were merged, and the regions with more than three significant SNPs were defined as high-confidence association loci. To define an eQTL hotspot, we narrowed each eQTL to 20 kb size according to up and downstream 10 kb of a leading SNP within an eQTL. We merged the narrowed eQTL if detected from different genes and overlapped. Finally, if an eQTL is associated with gene expression DMP in more than 44 genes (top 5%), we define this eQTL as a hotspot.

We estimated the dominance index (the ratio of dominance to additive effects, *d/a*) for each eQTL identified in the hybrid population, following the method described by (Zhou et al., 2019). For each eQTL, the median gene expression or MPH value associated with each allele of the lead SNP was calculated. Dominance (*d*) and additive effects were then derived using the formulas: *d* =F1 − mp and *a* =HP − mp, while F1 is the phenotypic value of the hybrid, mp is the mid-parent value, and HP is the value of the high parent. The level of dominance effect for each eQTL was determined as in our previous study (Liu et al., 2020): overdominance, *d/a >* 1.25; dominance, 0.25 *< d/a <*=1.25; additive, −0.25 *< d/a <*=0.25; recessive, −1.25 *< d/a <*=−0.25; and underdominance, *d/a <*=−1.25.

### Transcriptome-wide association study (TWAS)

We perform TWAS between hybrid gene expression and MPH of seedling traits using the Compressed Mixed Linear Model implemented in the package R GAPIT (Lipka et al., 2012; Zhang et al., 2010). Due to the format of the input file, the expression values were linearly transformed into values between 0 and 2 as in the previous study (Li et al., 2021). Similarly to eQTL mapping, the model utilized hidden factors and the first three PCs as covariates to control for population structure, and the kinship matrix was employed to control for genetic relatedness. The threshold for significant genes was set to 1.8e-6 (0.05/n, where *n* is the number of genes).

### SNP-based heritability and the genetic load

We defined SNP-based heritability 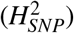 of gene expression as the proportion of total genetic variance explained by SNPs relative to the total variance in expression levels. For the hybrid population, we considered both additive and dominance genetic variance in estimating gene expression heritability. For the calculation, we employed 11.3 million SNPs to calculate the genetic relatedness matrix using the GCTA software (Yang et al., 2011; Zhu et al., 2015).

To estimate the genetic load in the 5-kb upstream regions of genes in inbred lines, we obtained Genomic Evolutionary Rate Profiling (GERP) scores from a previous study (Sun et al., 2023). SNP sites with GERP scores greater than zero were considered deleterious loci. We recoded genotypes as 0, 1, or 2, where 0 indicates a homozygous genotype matching that of sorghum (used as a proxy for the ancestral allele), 1 indicates a heterozygous genotype, and 2 represents the alternative homozygous genotype. For each SNP, we calculated the genetic load by multiplying the GERP score by the recoded genotype under an additive mode of inheritance. The total genetic load for the 5-kb upstream region of each gene was calculated by summing the individual SNP loads and then normalizing it by the region’s length (5 kb).

### Co-expression network construction and GO term enrichment test

We constructed the gene co-expression network using the WGCNA (v.1.73) (Langfelder and Horvath, 2008). For GO term analysis, we employed the HEMU online tool (Zhu et al., 2024), with a significance cutoff set at an FDR value of *<* 0.01.

### Clustering and population structure analyses

Population structure of the inbred lines was analyzed using ADMIXTURE (v.1.3.0) (Alexander et al., 2009). We evaluated values of *K* =2 to 10 to determine the optimal number of subpopulations, and *K* =6 was identified as the most appropriate division for our dataset (**Figure S1**). Inbred lines with membership probabilities ≥ 0.5 were assigned to their respective groups. Pairwise identity-by-state (IBS) distances among inbred lines were calculated using plink (v1.90b6.18) with the parameter `--ibs-matrix` (Purcell et al., 2007). The resulting IBS similarity matrix was converted into a genetic distance matrix (1 − IBS), which was then used to construct a neighbor-joining (NJ) phylogenetic tree with the ape package (Paradis and Schliep, 2019) in R (v4.3.0). Visualization was performed using a R package ggtree with a circular layout (Yu et al., 2017).

### ChIP–seq and DAP-seq data processing

Low-quality reads from the raw reads were filtered with fastp 0.20.0 (Chen et al., 2018) with default parameters. The remaining reads were aligned to the maize B73 AGP v4 reference genome (Jiao et al., 2017) using Bowtie v.2.2.8 (Langmead and Salzberg, 2012). Then we removed the duplicated reads using Picard tools (Toolkit, 2019). We called peaks using MACS2 (Zhang et al., 2008) with following parameters “–keep-dup 1 –nomodel -q 0.01 –shift -100 –extsize 200”.

### Luciferase activity assay

The coding sequence (CDS) of *ZmR1* was cloned downstream of the CaMV 35S promoter by linearizing the pRI101 vector with BamHI and SalI restriction enzymes. Similarly, a 2000 bp fragment upstream of the target gene’s start codon (ATG) was cloned into the pGreen II 0800 vector, linearized with KpnI and SalI. The recombinant constructs were introduced into *Agrobacterium tumefaciens* strain GV3101, and positive colonies were selected and cultured. Bacterial suspensions were adjusted to an OD_600_ of 0.8 prior to infiltration. The empty vector pRI101 and pRI101-ZmR1 were co-infiltrated into six-week-old tobacco (*Nicotiana benthamiana*) leaves along with the target gene constructs. After 48 hours of incubation, D-fluorescein potassium salt was applied, and fluorescence signals were captured using an in vivo imaging system.

### Phenotype collection and validation of ZmR1

Three over-expression lines of *ZmR1* from the previous study (Luo et al., 2021) were obtained. For seedling phenotype collection and validation, seeds of these over-expression lines, and wildtype Jing724 line were all cultured with soil on pots in a greenhouse at 50% relative humidity under an 18-h light/6-h dark photoperiod for 14 days. Then at 8:00 am on day 14, the individual plants were rinsed with sterile water and dried with sterile filter paper. Finally, lines with at least three growth-consistent individuals of each line were chosen for phenotype collection. The phenotypes, including mesocotyl length (ML), seedling length (SL), and root length (RL), were measured using rulers; the total length (TL) was the sum of SL, ML, and RL. Then, the fresh weight (FW) was measured for each individual, and all individual plants from the same line were put into a 60 °C oven for at least 48 hours until consistent dry weight data was obtained. For each line, average dry weight (ADW) was calculated from consistent dry weight divided by the number of individuals to measure FW. Then the fresh individual plants of each line were snap frozen with liquid nitrogen and stored at -80°C for RNA extraction. All of the lines used for seedling treatments were also planted for field-phenotype collection. In 2025, we grew these materials in two locations in China: Hai’nan, Sanya (18°1510N, 109°3043E), and Tai’an, Shandong (36°11’07”N, 117°08’05”E). The plant height and stalk stiffness were measured after flowering. Stalk stiffness (SS) of the fifth (bottom, BSS), sixth (middle, MSS), and seventh (top, TSS) internodes were measured using a digital stem strength tester (Model YYD-1B, Zhejiang Top Instrument Co., China). Measurements included at least six biological replicates, with each biological replicate assessed using three technical replicates.

### Determination of lignin content

The fifth internode of the individual plant at the silking stage was harvested from the Tai’an location to determine lignin content using a lignin assay kit (Solarbio, Beijing, China), according to the manufacturer’s instructions. Three biological replicates were prepared for each genotype, the wildtype Jing724 line and the overexpression lines. Briefly, samples were first oven-dried at 80 °C to constant weight, then ground and passed through a 40-mesh sieve. Precisely 5 mg of the processed sample was transferred to a 1.5 mL Eppendorf tube, followed by the addition of 500 µL of Reagent 1. The tubes were incubated in a 65 °C water bath for 30 minutes, then centrifuged at 8,000 × g for 5 minutes. The supernatant was discarded, and 500 µL of Reagent 2 was added. Samples were vortexed for 5 minutes, centrifuged again (8,000 × g, 5 minutes), and the supernatant was removed.

This washing step was repeated with 500 µL of Reagent 3, including vortexing for 5 minutes, centrifugation (8,000 × g, 5 minutes), and supernatant removal. Subsequently, 500 µL of Reagent 4 and 20 µL of Reagent 5 were added to each tube. A blank control was prepared in parallel by combining 500 µL of Reagent 4 and 20 µL of Reagent 5 in a separate tube without sample material. All tubes were incubated at 80 °C for 40 minutes. After cooling to room temperature, 500 µL of Reagent 6 was added to each tube. Following a final centrifugation at 8,000 × g for 10 minutes, 20 µL of the supernatant was mixed with 980 µL of glacial acetic acid. Absorbance was measured at 280 nm to determine lignin concentration.

### Phloroglucinol staining

At the silking stage, the fifth internode of the stem was harvested and manually sectioned into transverse slices approximately 200 µm thick using a stainless steel double-edged razor blade (Gillette). These sections were then mounted onto glass slides. We then prepared a 10% phloroglucinol ethanol solution by dissolving 1 g of phloroglucinol (GLPBIO) in 95% ethanol. In addition, a separate 1 M hydrochloric acid (HCl) solution was prepared. For staining, 40 µl of the 1 M HCl solution was applied to the stem section, immediately followed by 40 µl of the phloroglucinol ethanol solution. The reaction was proceeded for approximately 10 seconds and then gently blotted the excess liquid using absorbent paper. A coverslip was placed over the section, and the sample was promptly examined under a 4 *×* objective lens using an optical microscope (OLYMPUS BX51). Images were acquired using DPController software.

## Data and code availability

The raw sequence data reported in this paper have been deposited in the Genome Sequence Archive (Chen et al., 2021) in the National Genomics Data Center (Members et al., 2024), China National Center for Bioinformation, Beijing Institute of Genomics, Chinese Academy of Sciences (GSA: CRA023776) that are publicly accessible at https://ngdc.cncb.ac.cn/gsa. The code used for the analysis can be accessed from the GitHub repository (https://github.com/jyanglab/Maize_Expression_heterosis).

## Acknowledgements

This project was supported by Science and Technology Innovation 2030 — Major Project (2023ZD04068 to H.L.), National Key Research and Development Program of China (2022YFD1201700 to H.L.), and the National Natural Science Foundation of China (32072009, 32472176 to H.L. and 32101705 to X.Y.), and the Agriculture and Food Research Initiative grant number 2022-67013-36560 from the USDA National Institute of Food and Agriculture to J.Y.. We gratefully acknowledge Mingfei Sun, Yichen Hao, Yanjun Li, Jinxiao Zhang, Zixian Zhou, Yanxiao Lv, and Baiyue Zhang for their assistance in phenotype collection and RNA extraction. The authors thank Dr. James A. Birchler for his constructive discussions on functional validation and the anonymous reviewers for their valuable suggestions.

## Author contributions

J.Y. and H.L. designed this work. X.Y., M.Z., C.K., Z.T., and M.L. generated the data. G.X., X.Y., M.Z., Y.Q., P.L., J.R.-I., J.Y., and H.L. analyzed the data. J.Y., G.X., X.Y., and H.L. wrote the manuscript.

## Supporting Information

### Supporting Tables

**Table S1**. Hybrids and inbred lines used in this study. (https://github.com/GenXu1/Maize_Expression_heterosis/blob/main/02table/TableS1_sample_info.xlsx)

**Table S2**. Mid-parent heterosis (MPH) for seedling and adult traits. (https://github.com/GenXu1/Maize_Expression_heterosis/blob/main/02table/TableS2_MPH_of_seeding_adult_traits.xlsx)

**Table S3**. Summary of sequencing reads obtained from RNA-seq. (https://github.com/GenXu1/Maize_Expression_heterosis/blob/main/02table/TableS3_RNA-seq_reads.xlsx)

**Table S4**. Non-additive genes identified by expression data. (https://github.com/GenXu1/Maize_Expression_heterosis/blob/main/02table/TableS4_Non-additive_gene.xlsx)

**Table S5**. Gene Ontology enrichment analysis was performed using the genes in each module of the co-expression network constructed from the hybrid population. (https://github.com/GenXu1/Maize_Expression_heterosis/blob/main/02table/TableS5_Gene_Ontology_for_modules.xlsx)

**Table S6**. Genes involved in the anthocyanin, lignin, and benzoxazinoid pathways in module 19. (https://github.com/GenXu1/Maize_Expression_heterosis/blob/main/02table/TableS6_anthocyanin_ligin_Bx_in_M19.xlsx)

**Table S7**. eQTLs detected in inbred and hybrid populations and their association with expression dominance. (https://github.com/GenXu1/Maize_Expression_heterosis/blob/main/02table/TableS7_eQTL.xlsx)

**Table S8**. eQTLs for gene expression of *SBEIIa* (*Zm00001d003817*) in inbred and hybrid populations, as well as associations with expression dominance. (https://github.com/GenXu1/Maize_Expression_heterosis/blob/main/02table/TableS8_GWAS_for_SBEIIa_Zm00001d003817.xlsx)

**Table S9**. eQTL hotspots for expression dominance. (https://github.com/GenXu1/Maize_Expression_heterosis/blob/main/02table/TableS9_eQTL_hotspot_for_expression_dominance.xlsx)

**Table S10**. Transcriptome-wide association studies for seedling trait *per se* and their heterosis. (https://github.com/GenXu1/Maize_Expression_heterosis/tree/main/02table/TableS10_TWAS_results.xlsx)

### Supporting Figures

**Figure S1.**
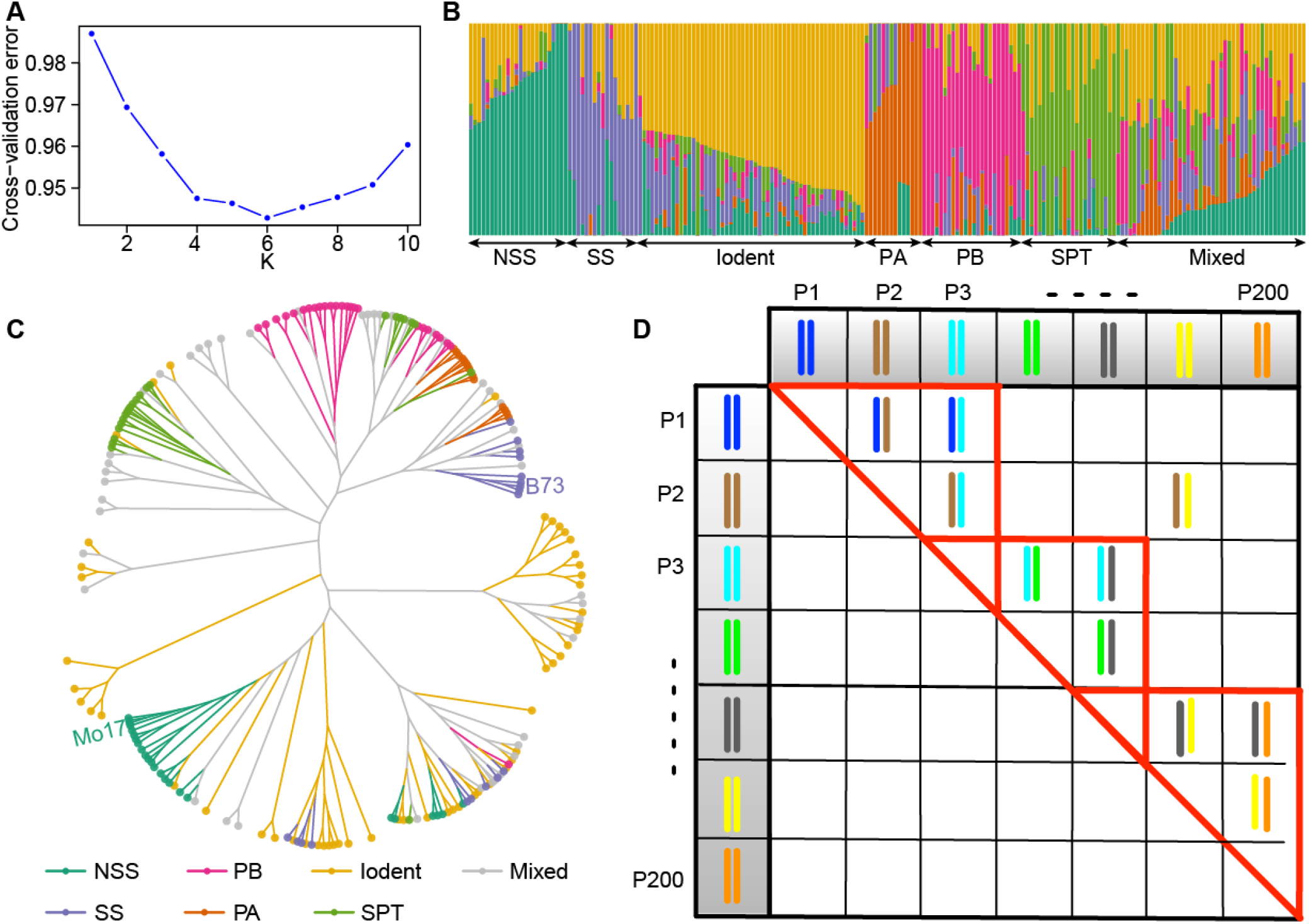
Population structure, phylogenetic relationships, and hybrid design of the maize inbred lines. (**A**) Cross-validation error plot from ADMIXTURE analysis used to determine the optimal number of subpopulations (*K* =6). (**B**) Population structure of the 203 maize inbred lines inferred by ADMIXTURE (*K* =6). Each vertical bar represents one inbred line, and colors indicate the estimated membership fractions of the six heterotic groups (NSS, SS, Iodent, PA, PB, SPT, and Mixed). SS, stiff stock; NSS, non-stiff stock; PA, group A germplasm derived from modern US hybrids; PB, group B germplasm derived from modern US hybrids; SPT, Sipingtou. (**C**) Neighbor-joining phylogenetic tree of the inbred lines constructed from genome-wide SNPs. Branch colors correspond to the heterotic groups defined in panel (**B**). (**D**) Schematic diagram illustrating the hybrid combinations derived from the parental lines shown in panel (**C**). In this design, around 200 diverse maize inbred lines were grouped into 10 blocks based on flowering time. Within each block, crosses were performed using a partial diallel mating design. Occasionally, crosses between blocks were also conducted.

**Figure S2.**
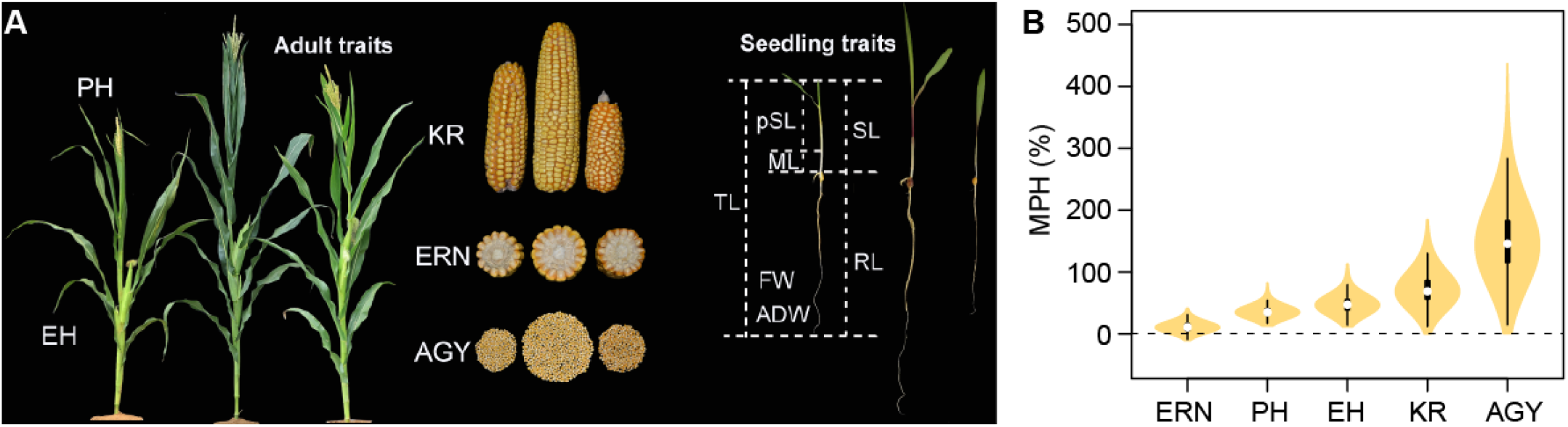
Data collection of adult and seeding traits. (**A**) The adult traits we collected in this study. Kernels were planted in the field to collect adult traits, including ear row number (ERN), plant height (PH), ear height (EH), kernels per row (KR), and average grain yield (AGY). Seedlings germinated from manually dissected embryos were used to collect seedling traits, encompassing fresh weight (FW), average dry weight (ADW), mesocotyl length (ML), root length (RL), seedling length (SL), partial seedling length (pSL), and total length (TL). Note that pSL = SL − ML, and TL = SL ====?RL. (**B**) Distribution of mid-parent heterosis (MPH) values for adult traits.

**Figure S3.**
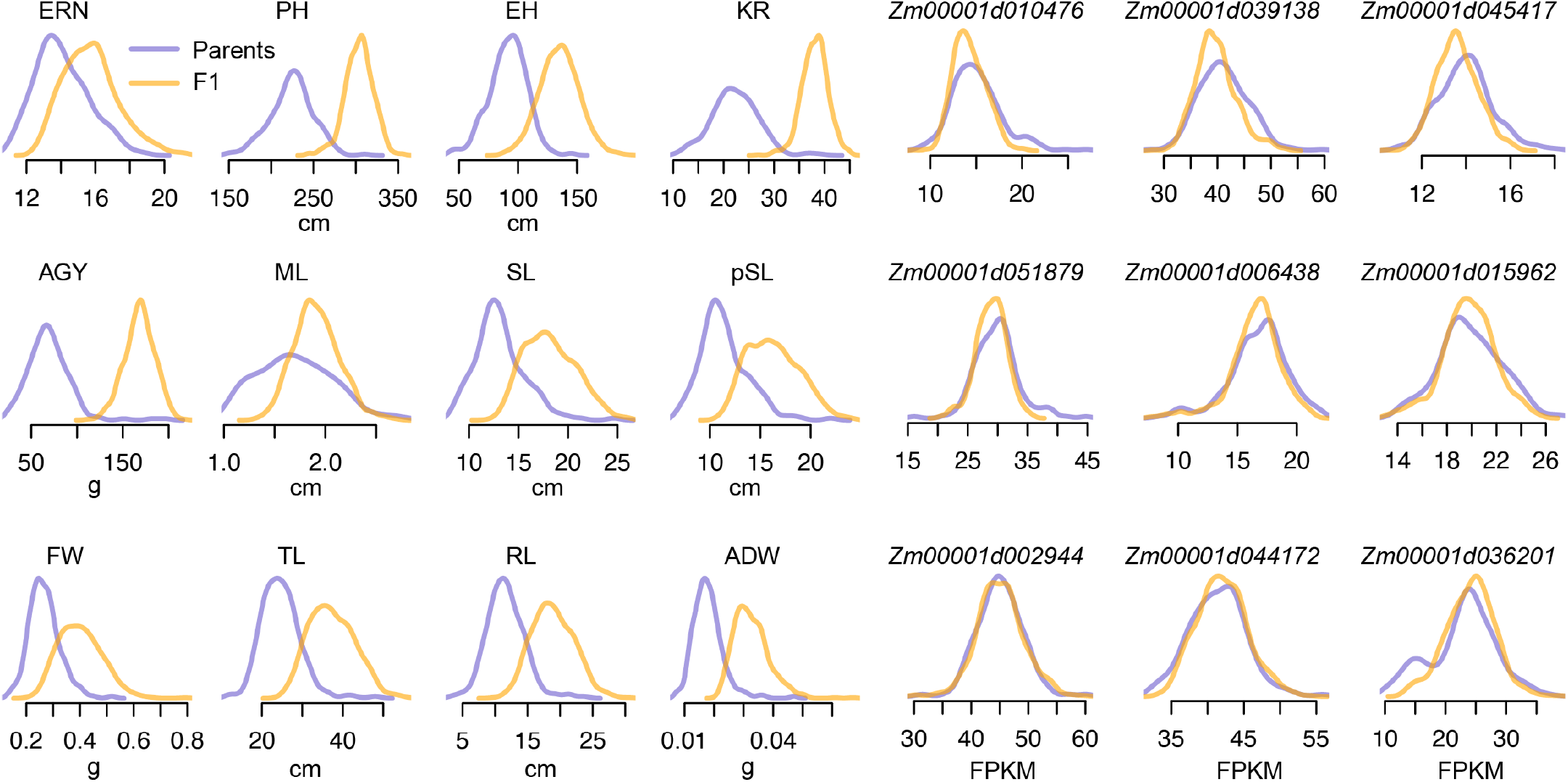
Distribution of adult, seedling, and molecular traits in inbreds and hybrids. The gene expression levels of nine housekeeping genes were shown for the molecular traits.

**Figure S4.**
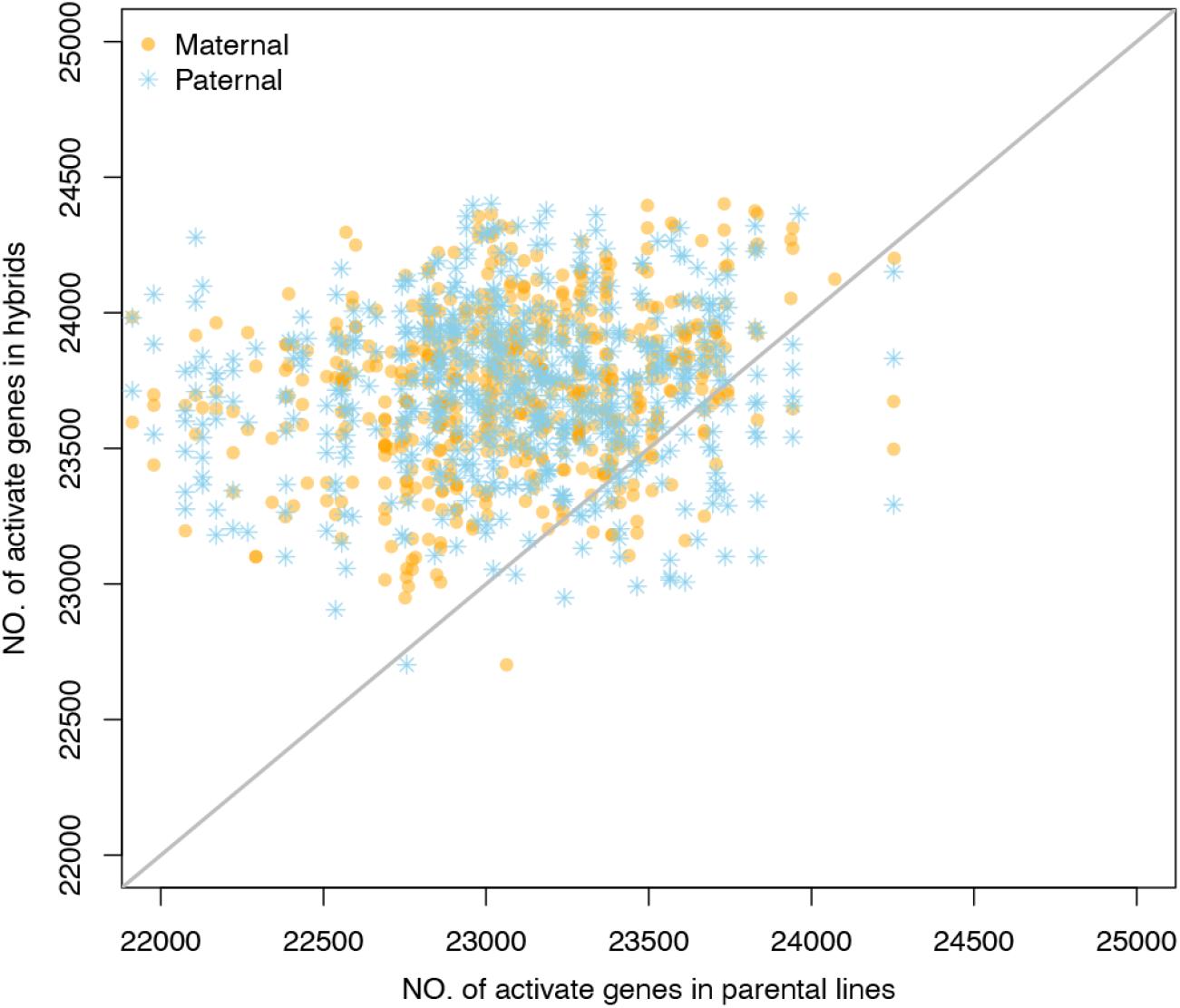
Number of expressed genes in hybrids and their parental lines.

**Figure S5.**
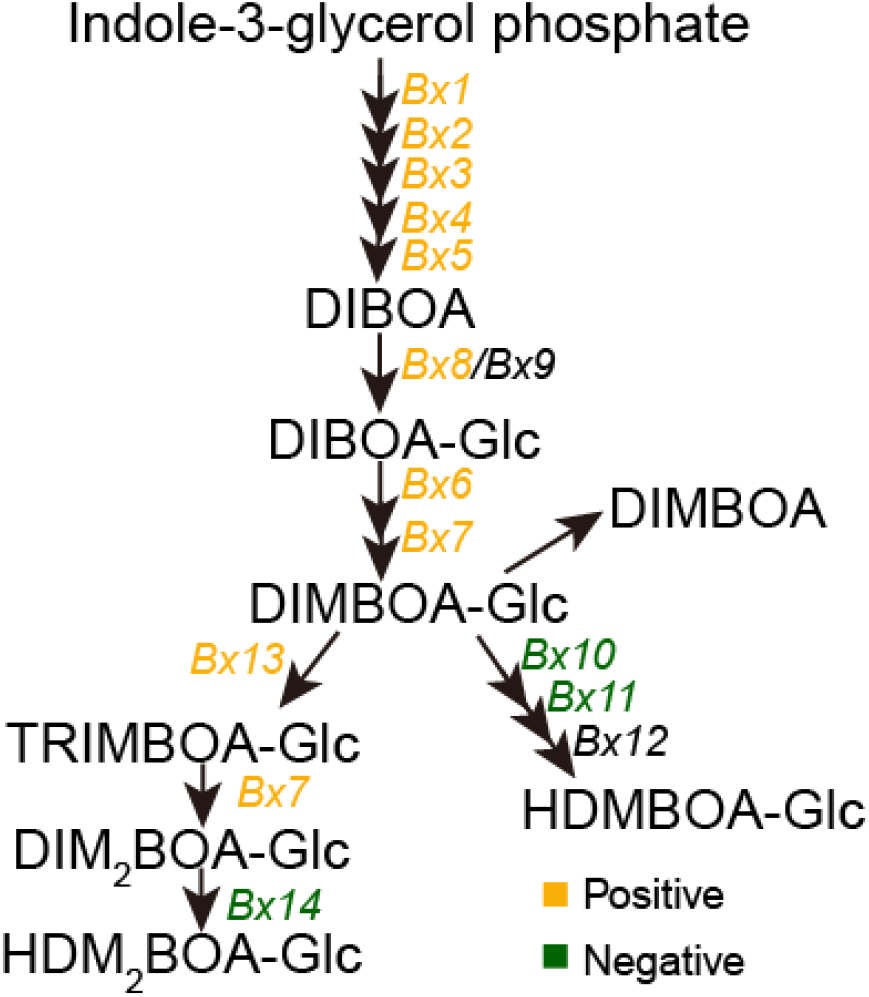
Benzoxazinoid metabolic pathway in maize. *Bx* genes highlighted in yellow indicate positive deviation from mid-parent (DMP) for gene expression, whereas those in green indicate negative DMP.

**Figure S6.**
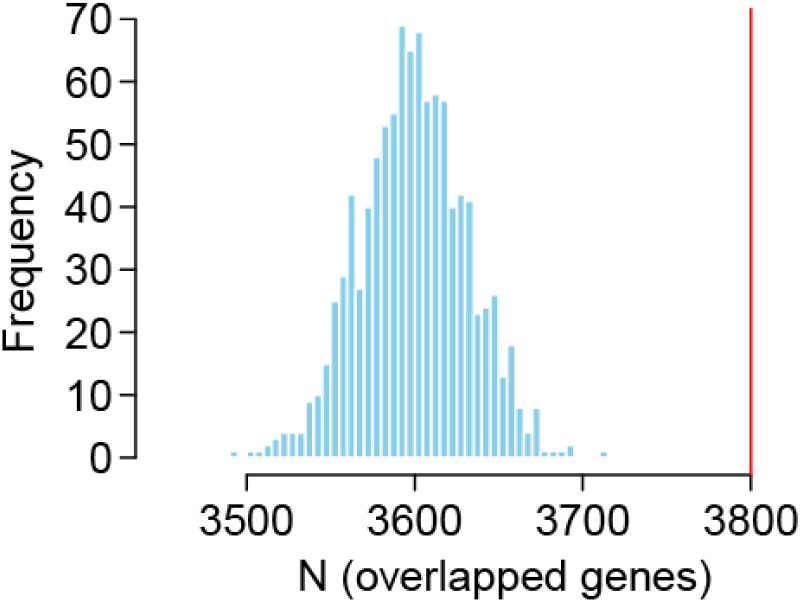
Comparison between non-acditive genes and syntenic genes. The number of syntenic genes overlapped with non-acditive genes. The red vertical lines indicate the observed value, and the histograms indicate 1000 permutation results using randomly selected expressed genes. The syntenic orthologs were determined by comparing maize with sorghum.

**Figure S7.**
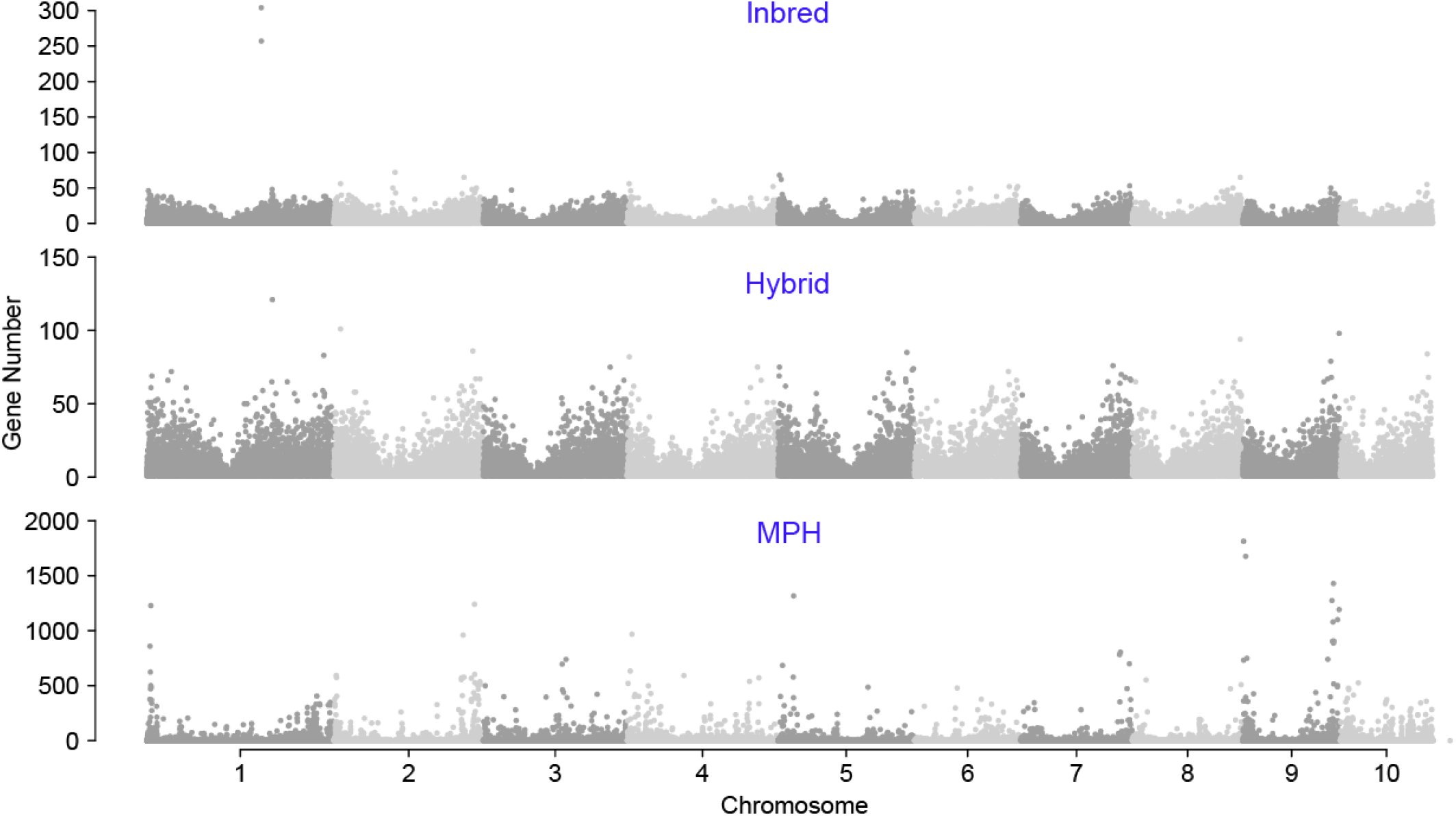
Distribution of eQTLs identified in inbred and hybrid populations, as well as those associated with expression dominance (DMP). The y-axis represents the number of genes regulated by eQTLs.

**Figure S8.**
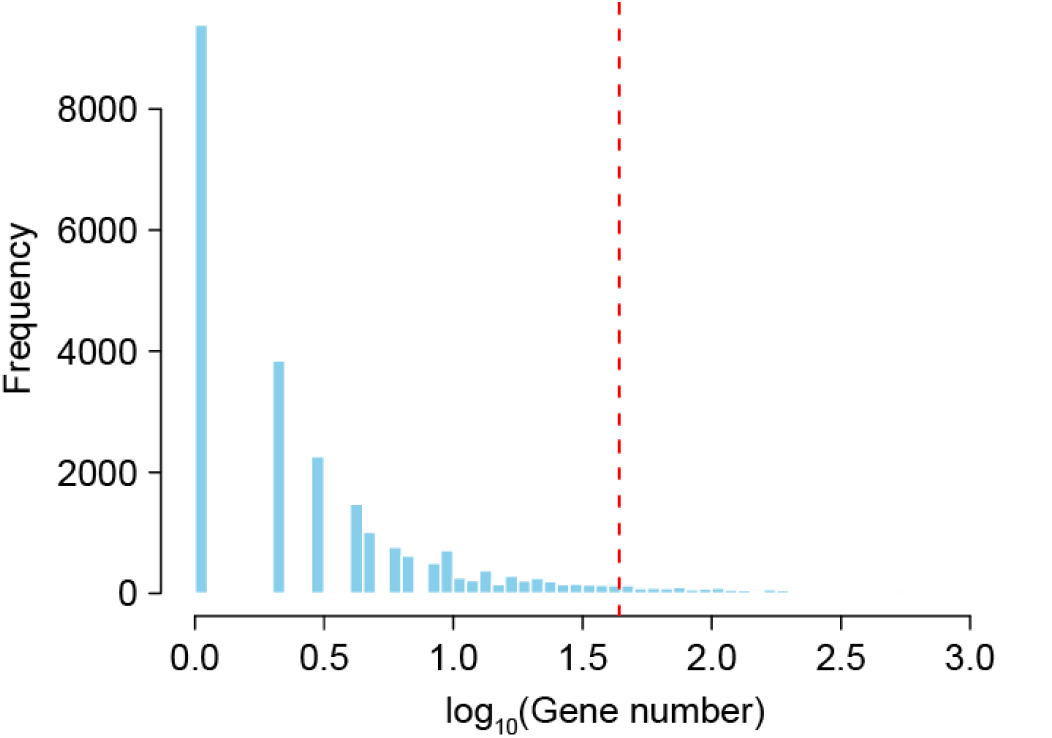
Distribution of gene number regulated by eQTLs for expression dominance. The red vertical line indicates the top 5% threshold.

**Figure S9.**
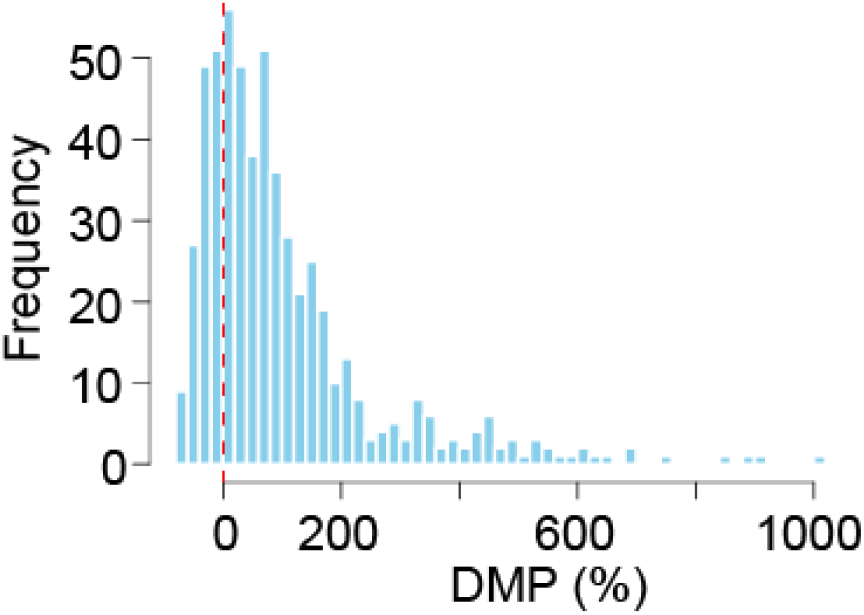
Distribution of deviation from mid-parent value (DMP) of *ZmR1* expression in hybrid population.

**Figure S10.**
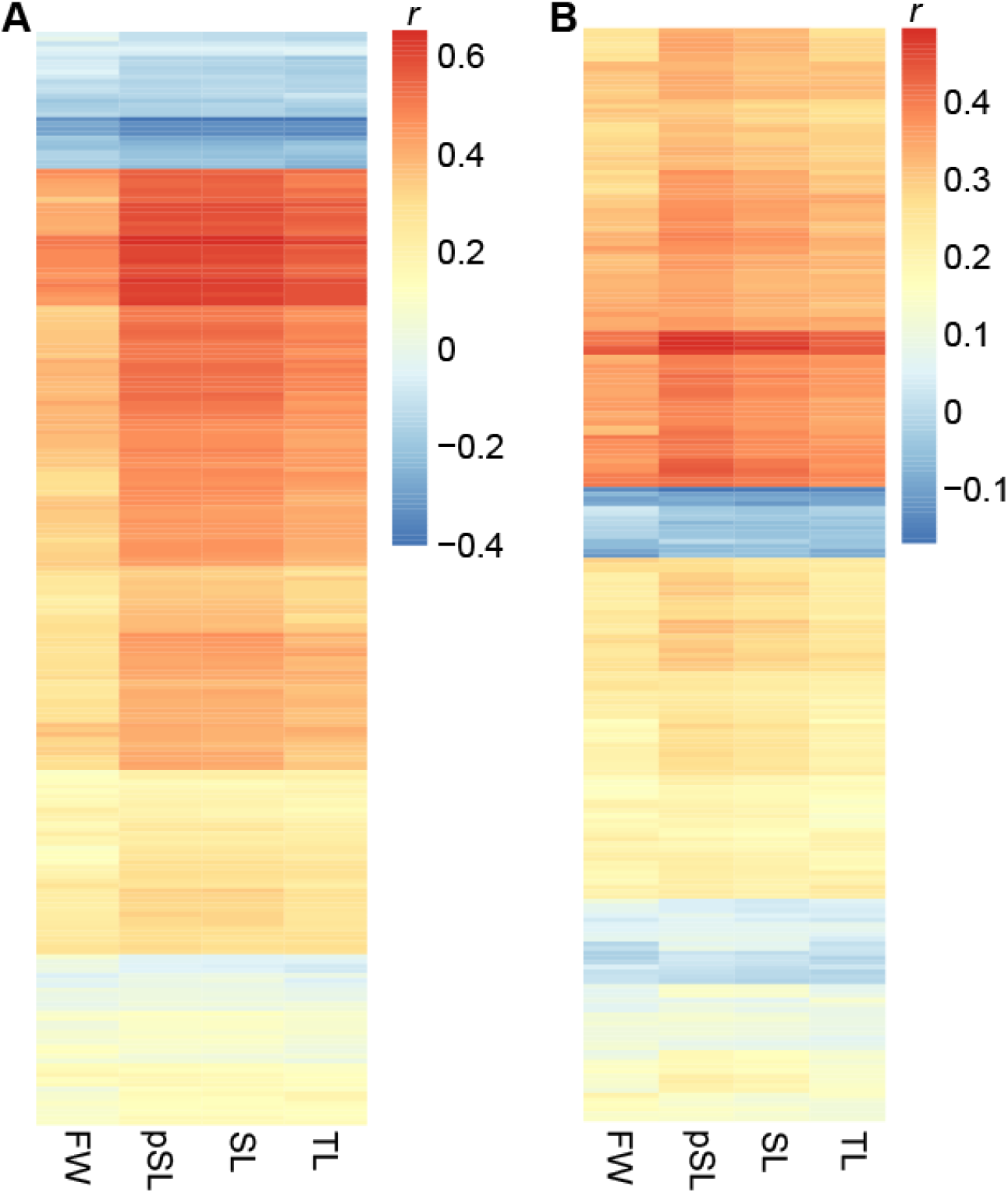
Correlation analysis between *ZmR1* target genes and seedling traits. (**A**) Correlations between the expression levels of *ZmR1* target genes and seedling trait values. (**B**) Correlations between the expression dominance (DMP) of *ZmR1* target genes and seedling trait heterosis values. FW, fresh weight; SL, seedling length; pSL, partial seedling length; TL, total length.

**Figure S11.**
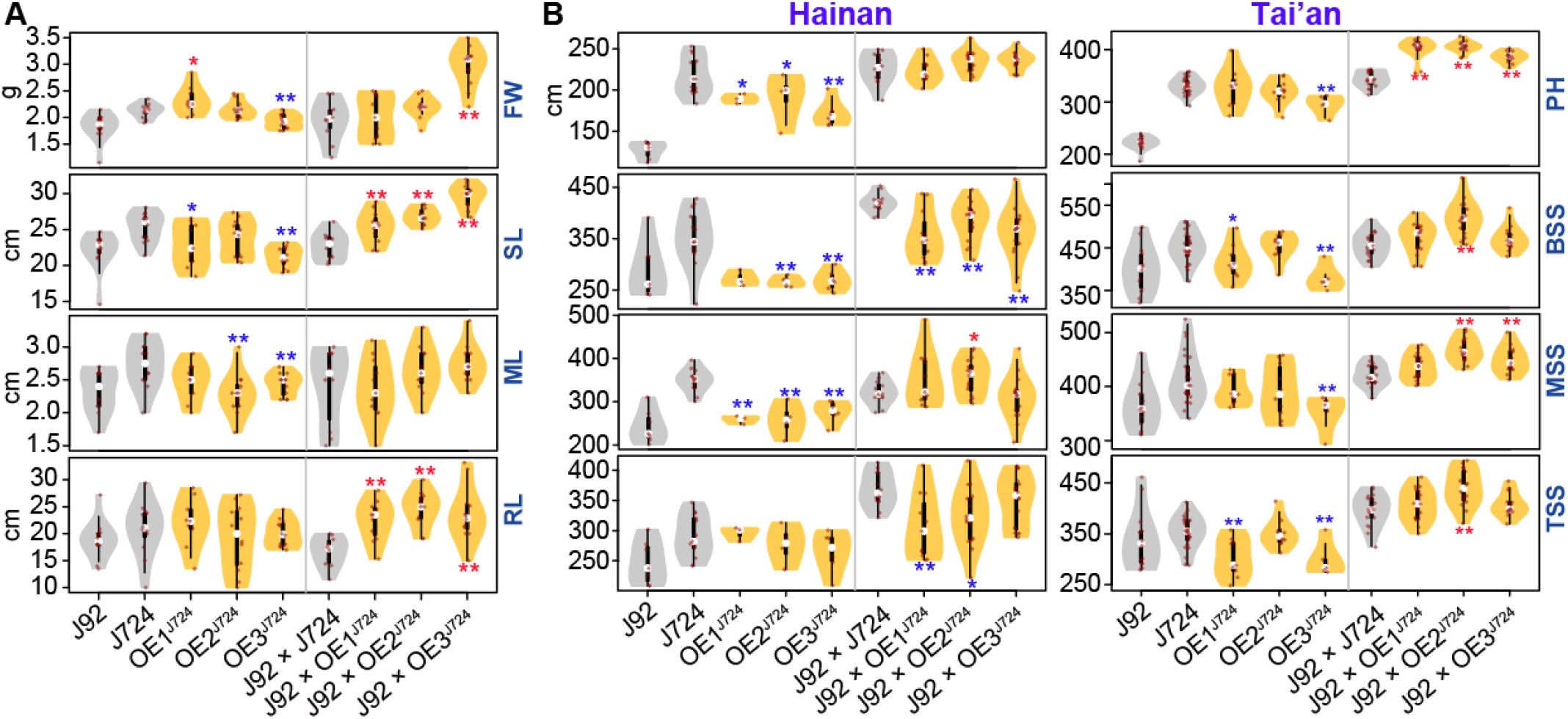
Phenotypic performance of *ZmR1* overexpression lines. Phenotypes of *ZmR1* overexpression lines as well as their hybrids, collected at seedling (**A**) and adult stages (**B**). FW, fresh weight; SL, seedling length; ML, mesocotyl length; RL, root length; PH, plant height; BSS, bottom stalk stiffness; MSS, middle stalk stiffness; TSS, top stalk stiffness. The adult traits were collected from two locations—Hainan and Tai’an, separately. Asterisks indicate significance levels between overexpression lines and their corresponding wildtypes, as determined by Wilcoxon’s test (*, *P* ≤ 0.05; **, *P* ≤ 0.01). Blue and red stars indicate overexpression lines exhibiting significant decreases or increases, respectively, relative to the wildtypes.

**Figure S12.**
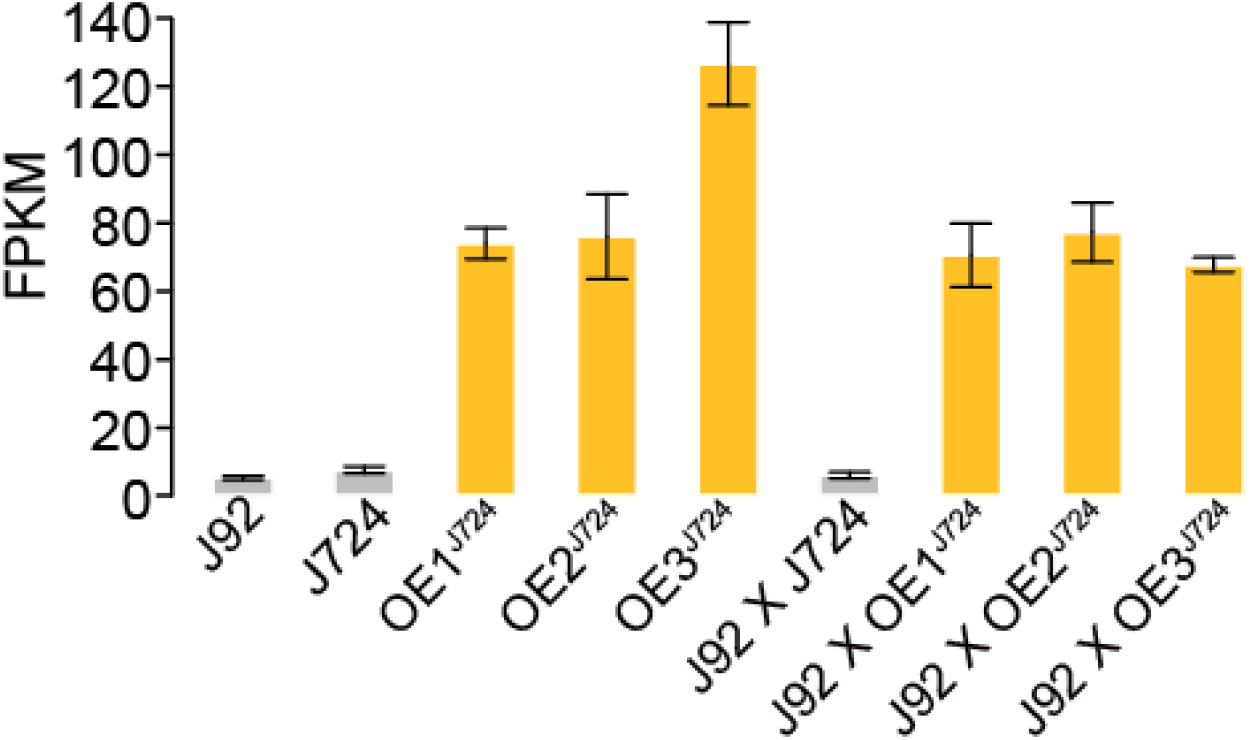
Expression levels of *ZmR1* in overexpression lines and their hybrids.

**Figure S13.**
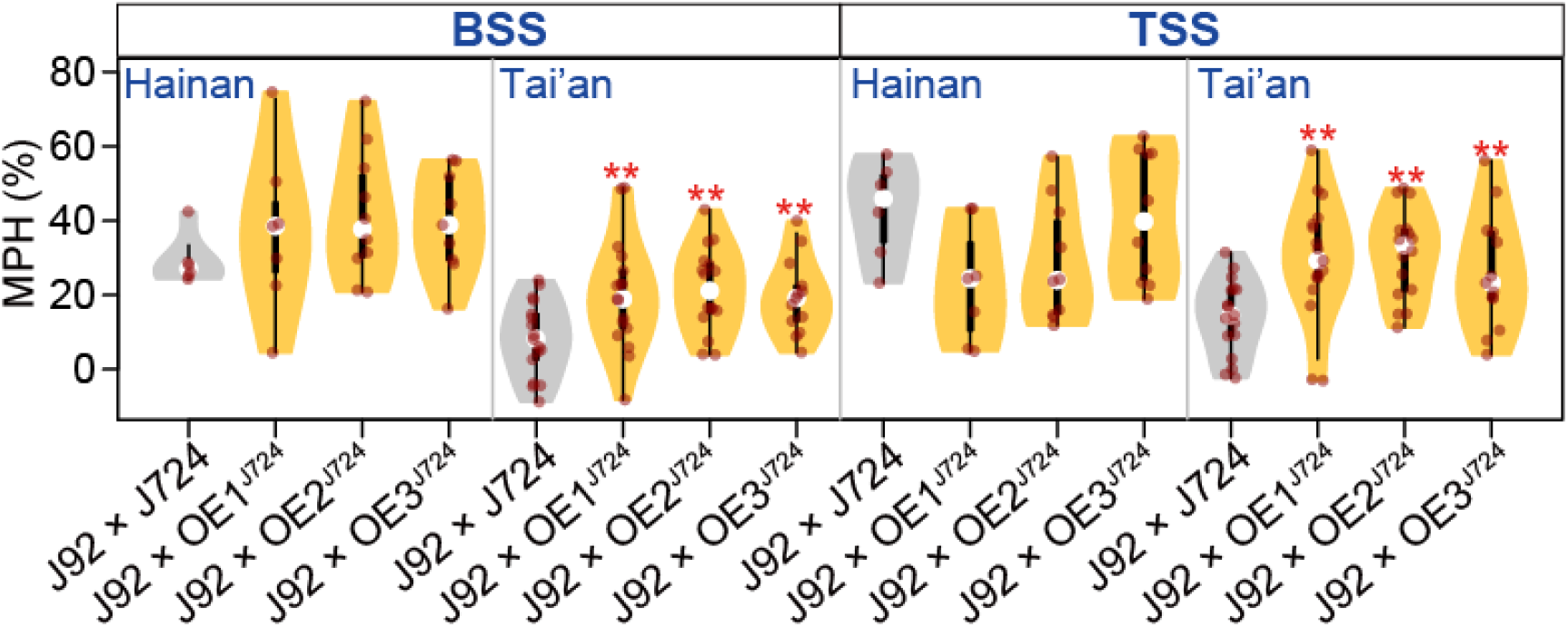
Heterosis of stalk stiffness in *ZmR1* overexpression lines. The stalk stiffness data was collected from two locations—Hai’nan and Tai’an, separately. TSS, top stalk stiffness; BSS, bottom stalk stiffness. Asterisks indicate significance levels between the overexpression lines and their corresponding wildtypes based on Wilcoxon’s test (*, *P* ≤ 0.05; **, *P* ≤ 0.01).

**Figure S14.**
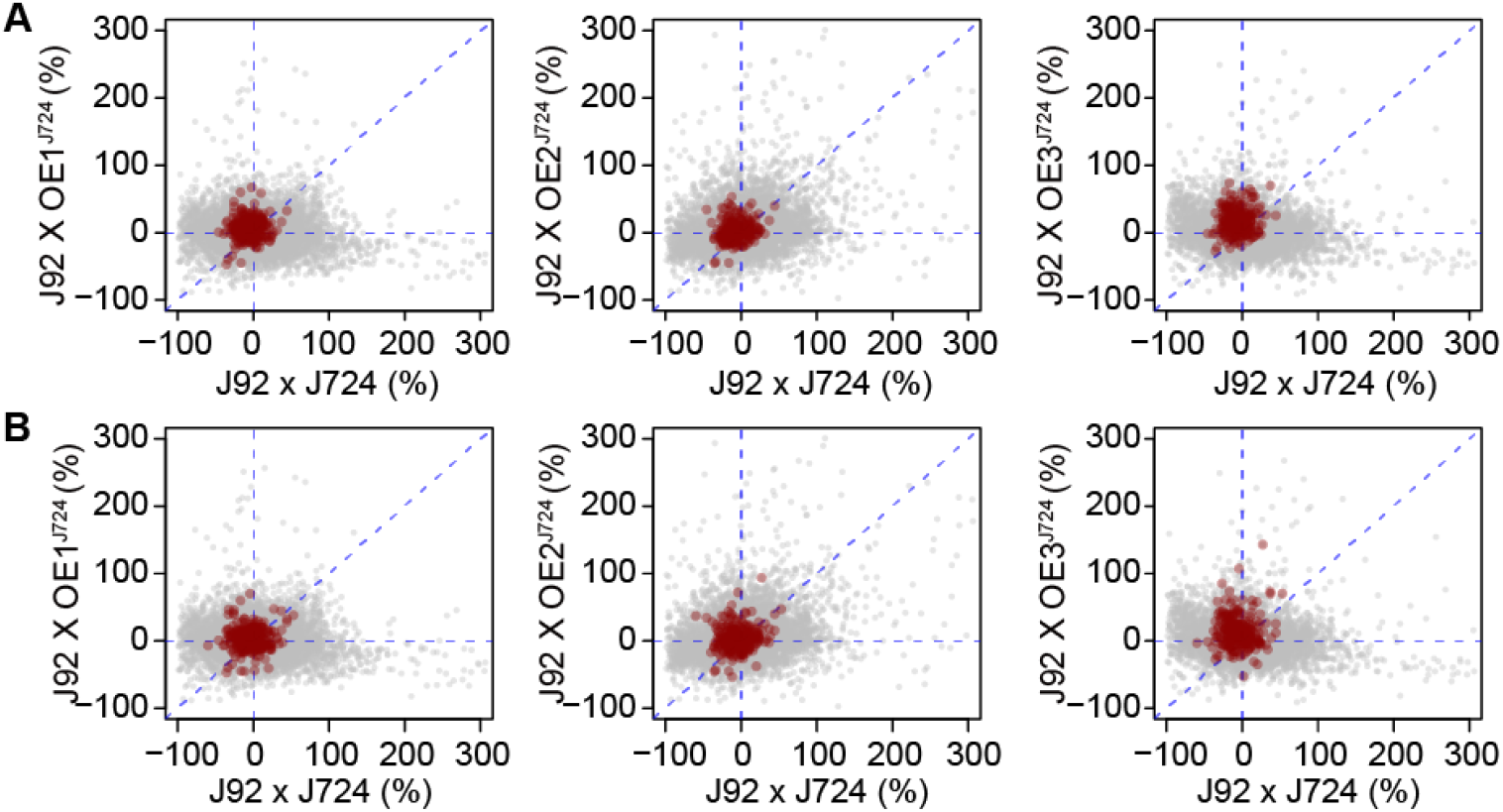
Comparison of expression DMP between hybrids derived from overexpression lines and their wildtype hybrids. Dark red dots represent *ZmR1* co-expressed genes in the inbred (**A**) and hybrid (**B**) populations.

